# Unraveling the role of MADS transcription factor complexes in apple tree dormancy using sequential DAP-seq

**DOI:** 10.1101/2021.02.21.431301

**Authors:** Vítor da Silveira Falavigna, Edouard Severing, Xuelei Lai, Joan Estevan, Isabelle Farrera, Véronique Hugouvieux, Luís Fernando Revers, Chloe Zubieta, George Coupland, Evelyne Costes, Fernando Andrés

## Abstract

**Background:** The effect of global warming on dormancy and flowering patterns of crop trees threatens world-wide fruit production and food security. In Rosaceous tree species, it is believed that a group of genes encoding MADS transcription factors (TFs) controls temperature-mediated dormancy cycle. These genes are similar to *SHORT VEGETATIVE PHASE* (*SVP*) from *Arabidopsis thaliana* and referred as *DORMANCY-ASSOCIATED MADS-BOX* (*DAM*) genes.

**Results:** By making use of apple tree (*Malus* x *domestica*) as a model for Rosaceous species, we have investigated the function of MADS TFs during the dormancy cycle. We found that MdDAM and other dormancy related MADS TFs form multimeric complexes with MdSVPa, and that MdSVPa is essential for the transcriptional complex activity. Then, for the first time in non-model plant species, we performed sequential DNA Affinity Purification sequencing (seq-DAP-seq) to define the genome-wide binding sites of these MADS TF complexes. Target genes associated with the binding sites were identified by combining seq-DAP-seq data with transcriptomics datasets obtained by the inducible glucocorticoid receptor expression system, and reanalyzing preexisting data related to dormancy cycle in apple trees.

**Conclusion:** We have determined a gene regulatory network formed by MdSVPa-containing complexes that regulate the dormancy cycle in apple trees in response to environmental cues. Key genes identified with our genomic approach and the elucidated regulatory relationships provide leads for breeding fruit trees better adapted to changing climate conditions. Moreover, we provide novel molecular evidence on the evolutionary functional segregation between DAM and SVP proteins in the Rosaceae family.

## Background

Temperate trees spread over geographical regions presenting wide seasonal environmental fluctuations throughout the year. In order to optimize their reproductive success, they adjust their growth and flowering cycles to these seasonal recurrent conditions. This plasticity is conferred by environment sensing mechanisms and signaling pathways that reprogram a population of pluripotent cells in specific tissues, called meristems, which act like stem cells in animals. The reduction of the photoperiod and the decrease of temperatures in winter serve as signals to induce growth cessation, the formation of winter buds to protect the meristematic tissues, and a meristematic state of rest called endodormancy (or winter dormancy), during which visible growth is inhibited. Endodormant buds can only recover their competence for growth after exposure to a certain period of low temperatures [1,2], which is species- and cultivar-dependent and termed as the chilling requirement (CR). The satisfaction of the CR leads to a vast genetic reprogramming of the shoot apical meristem (SAM) that undergoes an ecodormant phase [3–7], i.e. competence to resume growth if experiencing a sufficient amount of warm temperatures (so-called heat requirement or HR). Therefore, in springtime, when climatic conditions are favorable for reproduction, budbreak and flowering take place. The timing of budbreak and flowering is dependent on yearly cycles of temperature oscillations and thus highly susceptible to climate change. The detrimental effects of global warming on agriculture are already visible [8]. In temperate fruit trees, warmer winters cause non-synchronized and defective flowering due to insufficient chilling accumulation, whereas higher temperatures registered in spring lead to early flowering and, consequently, fruit damage by spring frost [9].

The Rosaceae is the third most economically important plant family in temperate regions and responsible for the major part of the total worldwide-consumed fruits [10]. Iconic Rosaceous species such as apple (*Malus domestica*), peach (*Prunus persica*) and sweet cherry (*Prunus avium*) are characterized by high CRs. This feature limits the range of latitudes at which they can be productively grown and makes them highly susceptible to global warming. However, despite the importance of a well-adjusted dormancy cycle for flowering timing and fruit production, the understanding on how this process is controlled in Rosaceae and other significant crops is still in its infancy. It is believed that a group of genes encoding MADS transcription factors (TF), named for founding members <u>M</u>CM1 (*Saccharomyces cerevisiae*), <u>A</u>GAMOUS (*Arabidopsis thaliana*), <u>D</u>EFICIENS (*Antirrhinum majus*) and Serum response factor (*Homo sapiens*), are major regulators of the dormancy cycle in many Rosaceous fruit trees [11–13]. These genes are referred as *DORMANCY-ASSOCIATED MADS-BOX* (*DAM*) and were first identified because of their genetic association with the non-dormant phenotype of the *evg* mutant of peach [14]. *DAM* genes belong to the MIKC^C^ type of MADS TF genes and are similar to *SHORT VEGETATIVE PHASE* (*SVP*) from *Arabidopsis thaliana* [15]. Further genetic studies have identified *DAM* genes within QTLs related to dormancy cycle in diverse fruit tree species, such as apple [16], pear [17], peach [18], almond *[Prunus dulcis* (Miller) D. A. Webb] [19], apricot (*Prunus armeniaca* L.) [20] and sweet cherry [21]. Notably, the expression of *DAM* genes from these and many other fruit tree species display distinct seasonal patterns [11]. The mRNA transcription of a subgroup of *DAM* genes (referred as “pattern #2” by [11]) is induced by low temperatures and shows a maximum peak of expression in the middle of the winter. This peak of expression has been associated with the physiological establishment and maintenance of endodormancy [22]. Characteristically, the expression of “pattern #2” *DAM* genes progressively decreases after a long exposure to cold and reaches its minimum around the time when CR is satisfied and endodormancy is released [3,11]. The function of some *DAM* genes showing this particular pattern of expression has been studied in transgenic trees. Ectopic expression of the Japanese apricot (*Prunus mume* Sieb. et Zucc.) *PmDAM6* gene in poplar induced growth cessation, bud set and bud endodormancy [22]. The silencing of the apple *MdDAM1* expression in transgenic apple trees led to the opposite phenotype, as plants showed a non-dormant and constant growing phenotype [4]. A similar phenotype was observed in transgenic apple trees in which three *MdDAM* genes (*MdDAM1, MdDAM4* and *MdDAMb*) and two *MdSVP* genes (*MdSVPa* and *MdSVPb*) were simultaneously downregulated using a synthetic RNA interference molecule [23]. Conversely, the overexpression of *MdDAMb* and *MdSVPa*, which patterns of expression are different to those described for *MdDAM1* and *PmDAM6* [22,24], caused growth inhibition and delayed budbreak, but did not affect growth cessation and endodormancy induction [25]. The role of *SVP* ortholog genes (*SVP*-like) in repressing budbreak has been further demonstrated in the forestry tree model species hybrid aspen [26]. The functional characterization of *DAM* and *SVP*-like genes suggests the existence of one or more temperature-mediated gene regulatory networks (GRNs) controlling the dormancy cycle in which these genes play a central role. Deciphering these GRNs is essential to better understand how budbreak and flowering is timely modulated in fruit trees, and would provide means to breed for early- and late-flowering cultivars better adapted to climate change.

In Arabidopsis, SVP inhibits floral transition by affecting GRNs that integrate environmental cues and endogenous hormonal signals [27]. SVP directly binds to and represses the expression of the floral integrator genes *FLOWERING LOCUS T* (*FT*) and *SUPPRESSOR OF OVEREXPRESSION OF CONSTANS 1* (*SOC1*), besides controlling the expression of several genes related to gibberellin (GA) biosynthesis and signaling [28–30]. Remarkably, as many other MADS TFs in flowering plants, SVP forms complexes of dimers or tetramers that bind to specific DNA sequences termed CArG-box (CC(A/T)_6_GG) [31,32]. The oligomeric composition of these complexes defines the genomic regions to which they bind, allowing multiple transcriptional responses by combination of distinct TFs [33]. In Arabidopsis, SVP physically interacts with FLOWERING LOCUS C (FLC) to form a transcriptional repressive complex that inhibits floral transition [29]. FLC is a MADS TF that represses flowering until the plant is exposed to a prolonged period of low winter temperatures in a process known as vernalization [34]. In Brassicaceae species, it has been shown that vernalization induces the progressive *FLC* repression by an epigenetic mechanism that involves the accumulation of trimethylation of lysine 27 on histone 3 (H3K27me3) [35]. By combining single and double *SVP* and *FLC* mutants together with transcriptomics and chromatin immunoprecipitation coupled to next-generation sequencing (ChIP-seq) studies, it has been possible to identify direct transcriptional targets of SVP, FLC and the SVP–FLC complex (including *FT*, *SOC1* and GA-related genes) [27,36,37]. Besides FLC, SVP form complexes with many other MADS TFs to modulate transcriptional networks involved in the thermosensory flowering pathway and floral development [27,38–40]. The identification of these complexes and the genome-wide binding sites bound by the studied MADS TFs have enormously contributed to decipher their individual and cooperative developmental roles in the model plant species Arabidopsis [27,36,41–43].

The molecular mechanism underlying dormancy cycle control by temperature in trees is believed to be analogous to that described for the vernalization pathway in Arabidopsis [11,44]. This notion is mainly supported by the above described cold-mediated transcriptional regulation of *DAM* genes, which resembles that of *FLC* in Arabidopsis, and by the similar role of *SVP*-like genes in repressing budbreak date and flowering-time in fruit trees and Arabidopsis, respectively [25,26,45]. Furthermore, the function of FLC-like TFs has been recently related to tree dormancy control [46–48]. However, there are no proofs yet on the existence of multimeric complexes formed by MADS TFs, such as DAM and SVP-like, that govern transcriptional networks controlling the dormancy cycle in trees. Here, we have made use of a recently developed technology called sequential DNA Affinity Purification sequencing (seq-DAP-seq) to gain knowledge on the function of apple MADS TFs. Seq-DAP-seq is based on a sequential protein immunoprecipitation with multiple tags to isolate heteromeric MADS TF complexes followed by DNA affinity purification [43]. By combining protein–protein interaction assays, seq-DAP-seq and transcriptomics, we isolated MADS TF complexes and defined their transcriptional genome-wide targets. Our results show that apple DAM-, FLC- and SVP-like complexes operate in GRNs, which integrate environmental and hormonal signaling pathways to regulate dormancy cycle progression. Moreover, we propose the existence of an evolutionary functional segregation between DAM and SVP-like TFs. Whereas SVP flowering-related functions are conserved between phylogenetically distant species, DAM TFs have diverged to play specific dormancy roles in Rosaceous tree species.

### Results

#### MdSVPa *and* MdSVPb *but not* MdDAM-like *genes complement the early-flowering phenotype of Arabidopsis* svp-41

Previous phylogenetic analysis classified the *DAM* genes as related to Arabidopsis *SVP*, therefore the *DAM* genes are commonly referred to as *SVP*-like genes [15]. However, this classification may be misleading because a broader phylogenetic analysis recently showed that these genomes carry two groups of *SVP*-related genes: a cluster solely composed of Rosaceous *DAM* genes and another well-defined cluster formed by Arabidopsis *SVP* and *SVP*-like genes from Rosaceous species [11]. This classification will be used to describe the genes of this study. Apple contains five *DAM*-like and two *SVP*-like genes. The CDS of four out of five *DAM*-like genes and both *SVP*-like genes was successfully amplified and cloned. To evaluate their functional homology to *SVP*, apple *DAM-* and *SVP*-like genes were misexpressed in an Arabidopsis mutant lacking *SVP* function (*svp-41*), which exhibits an early-flowering phenotype [45]. For this purpose, 3 kb of the *SVP* promoter was used to drive transgene expression. The Arabidopsis *SVP* gene was used as a positive control [49], and the Venus fluorescent protein was used to monitor the temporal and spatial expression pattern conferred by this promoter. T1 transformants were selected by BASTA application and scored for flowering time based on the number of rosette leaves at bolting (Supplemental Figure S1). Lines that showed a Mendelian segregation (3:1) in the next generation were followed, and homozygous single-copy T3 lines from independent T1 lines were scored for three flowering-time traits under long-day (LD) conditions. As expected, plants carrying Venus were not able to restore the WT phenotype of *svp-41*. Confocal microscopy of these plants revealed that the *SVP* promoter used in this study was able to correctly guide the expression of the transgenes throughout expanded leaves and the SAM (Supplemental Figure S2) [28,45,50]. Plants expressing *SVP*, *MdSVPa* and *MdSVPb* were reproducibly late flowering under LDs in comparison to *svp-41*. These plants produced three to seven more leaves prior to flowering (Figure 1A); bolted to form an inflorescence 6–9 days later (Figure 1B); and flowered 3–7 days later than *Venus svp-41* plants (Figures 1C and 1E). Lines showing an intermediate phenotype contained lower transgene mRNA levels than their counterparts (Figure 1D). Lines carrying *MdDAM*-like genes showed transgene mRNA levels similar to those observed in plants expressing *SVP*, *MdSVPa* and *MdSVPb*. However, they could not complement any of the *svp-41* floral phenotypes. No statistical significance was observed in any of the flowering traits analyzed among plants expressing *SVP*, *MdSVPa* or *MdSVPb*, which suggests that apple *SVP*-like genes share functional homology to Arabidopsis *SVP*; whereas *MdDAM* genes do not. These data suggest an evolutionary diversification between *DAM*- and *SVP*-like genes within the Rosaceae.

**Figure 1.**
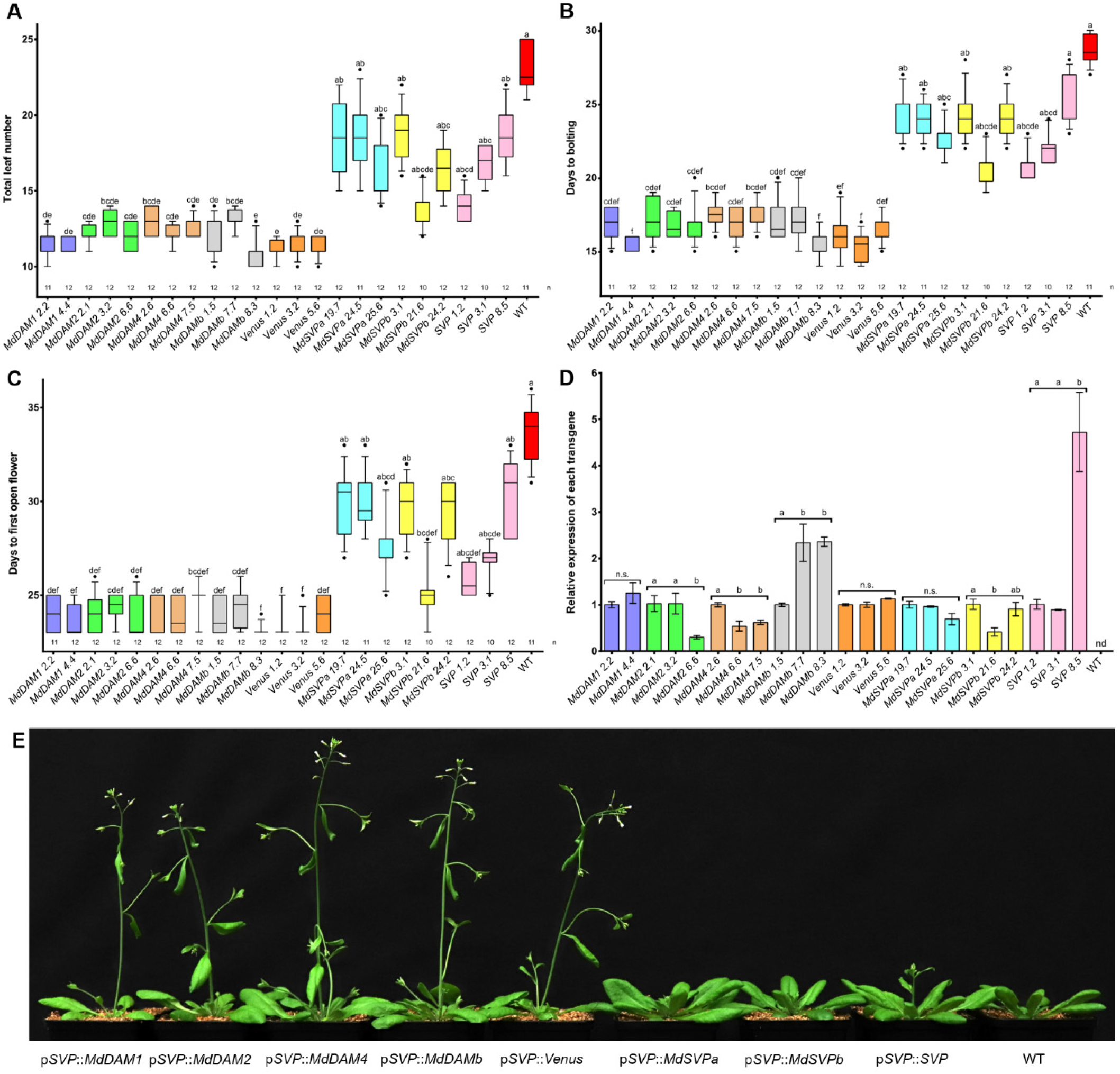
Complementation assay of the early-flowering phenotype of Arabidopsis *svp-41* mutant under LDs. **A)** Total leaf number (including both cauline and rosette leaves) was scored prior to flowering. **B)** Number of days from germination to bolting (elongation of the first internode around 0.5 cm). **C)** Number of days from germination to the opening of the first flower. In **A** to **C**, the box extends from the 25th to 75th percentiles, the line in the middle is plotted at the median, and the whiskers are drawn down to the 10th and up to the 90th percentile. The outliers below and above the whiskers are drawn as individual points. Kruskal-Wallis one-way ANOVA followed by Dunn’s test was used for the statistical tests. Letters shared in common between the genotypes indicate no significant differences (for P ≤ 0.05). **D)** Relative expression of the different transgenes after 7 LDs. Gene expression from three independent biological replicates ± SEM is shown relative to the reference gene *PP2A*, with the first genotype of each transgene set to 1. Statistical analysis was done using one-way ANOVA followed by Tukey’s multiple comparison test or t test for *MdDAM1* comparisons. Letters shared in common between the genotypes indicate no significant differences (for P ≤ 0.05). n.s = nonsignificant; nd = non-detected. **E)** Plants representing intermediate phenotypes of each genotype grown under LDs for 25 days.

#### MdSVPa interacts with several apple MADS TFs related to dormancy-specific phases

Dimerization is a key feature of MADS TFs, and their combination in protein complexes defines their function by conferring target specificity [51,52]. Thus, whether apple DAM- and SVP-like proteins form complexes was tested. Additionally, due to the well-characterized SVP–FLC module in Arabidopsis [36], the apple FLC-like (MD09G1009100) protein was added to these interaction tests [46]. In a first attempt, full-length coding sequences were used to screen protein–protein interactions using yeast two-hybrid (Supplemental Figure S3A). Interactions between MdSVPa and MdDAMb, MdSVPa and MdFLC, and the MdSVPa homodimer were detected. However, MdDAM1, MdDAM2 and MdDAM4 showed strong autoactivation of yeast reporter genes when fused to the DNA-binding domain. It is well-described that the autoactivation generated by MADS proteins can be abolished by removing their C-terminal region [53,54]. Truncated versions missing their C terminus were generated for all seven proteins, and interactions were tested in a reciprocal yeast two-hybrid assay unless a positive interaction was detected beforehand (Figure 2A). MdDAM1 and MdDAM4 interacted with both MdSVP-like proteins, and a positive interaction was also identified between MdSVPa and MdSVPb. MdDAM4 and MdSVPb also showed the ability to form homodimers. All tested proteins were able to interact with MdSVPa, except MdDAM2 that did not interact with any apple DAM-, SVP- or FLC-like protein. To test whether these interactions also occur *in planta*, coimmunoprecipitation assays were carried out in agroinfiltrated *Nicotiana benthamiana* leaves (Figure 2B). For this assay, full-length coding sequences were tagged with N-terminal GFP or 5xMyC. Controls were used to ensure that the MADS TFs did not recognize and bind to GFP- or MyC-tag (Venus fused to a nuclear localization signal (NLS) and AtHB33-MyC, respectively). All protein–protein interactions observed in the yeast two-hybrid assay were successfully validated, demonstrating the existence of a physical interaction network among apple DAM-, SVP- and FLC-like proteins (Figure 2C). These analyses revealed MdSVPa as a central hub among these interacting proteins. The subcellular localization of the proteins was determined in agroinfiltrated tobacco leaves with NLS-Venus as a control (Supplemental Figure S4). All apple proteins were localized in the nucleus, providing further evidence that the protein–protein interactions here identified may have a transcriptional regulatory role.

**Figure 2.**
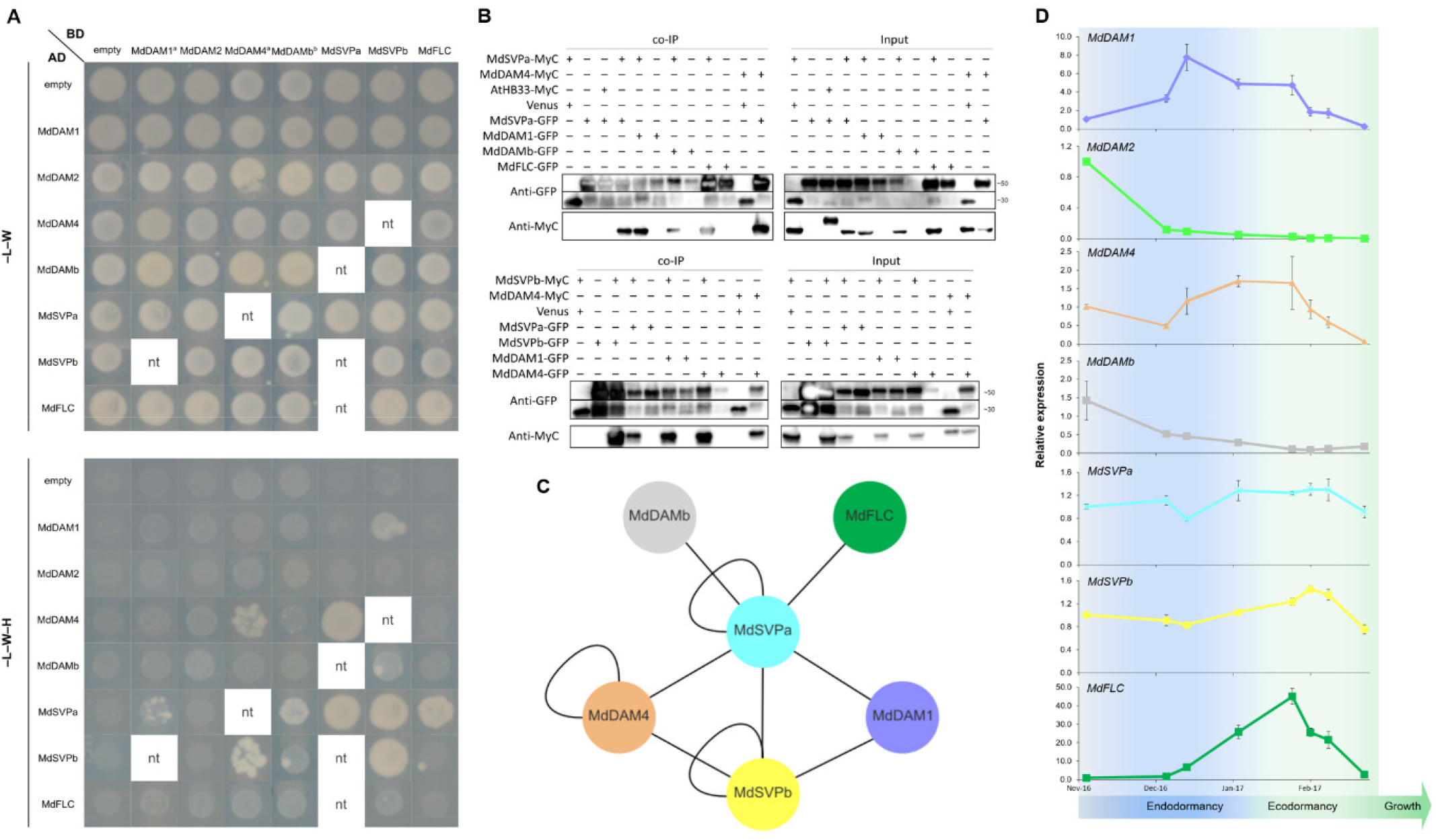
Protein interaction and gene expression analysis of genes encoding apple DAM-, SVP- and FLC-like proteins. **A)** Yeast two-hybrid assay of truncated apple MADS proteins. Interactions were reciprocally tested unless a positive interaction was identified beforehand. In this case, the other direction was not tested (blank spaces with nt – not tested). The negative controls are representative pictures of several independent controls. ^a^MdDAM1 and MdDAM4 fused to the DNA-binding domain were evaluated in –L–W–H media supplemented with 0.5 mM 3-AT. ^b^The interaction between MdSVPa and MdDAMb was observed using full-length protein versions. See Supplemental Figure S3 for further information. **B)** *In planta* co-immunoprecipitation of protein interactions detected by yeast two-hybrid. MdDAM4, MdSVPa and MdSVPb were translationally fused to MyC, whereas MdDAM1, MdDAM4, MdDAMb, MdSVPa, MdSVPb and MdFLC were translationally fused to GFP. Protein– protein interactions were tested in pairs by agroinfiltration of tobacco leaves. The input was composed of total proteins recovered before immunoprecipitation. GFP-fused proteins were immunoprecipitated using anti-GFP nanobody (VHH) beads and immunoblotted using anti-MyC or anti-GFP antibody. NLS-Venus was used to demonstrate that the MADS proteins do not bind to GFP, whereas AtHB33-MyC (AT1G75240) was used for the same purpose but for the MyC-tag. The anti-GFP blot was split in two due to differences in protein size between NLS-Venus (around 30 kDa) and the GFP fusions (around 50 kDa). **C)** Summary of the protein–protein interactions identified in this work. **D)** Relative expression of apple *DAM-, SVP-* and *FLC*-like genes during the dormancy cycle. Gene expression from three independent biological replicates ± SEM is shown relative to the reference genes *MdMDH* and *MdWD40*, with the first sampling point of each gene set to 1. Endo- to ecodormancy transition was determined by forcing tests.

In the light of these results, we tested whether the presence of MdSVPa could reveal a flowering phenotype attributable to MdDAM. The *MdSVPa 24.4 svp-41* line was crossed to *MdDAM1 4.4 svp-41, MdDAM2 3.2 svp-41, MdDAM4 6.2 svp-41, MdDAMb 1.5 svp-41* and *Venus 1.2 svp-41*, and the flowering time of F1 plants were compared with each parent (Supplemental Figure S5). No differences in flowering traits were observed in F1 lines misexpressing both transgenes versus *MdSVPa*. These results reinforced the notion that MdDAM and MdSVP TFs play different functions, at least in the context of the flowering-time control of Arabidopsis.

Next, we investigated the function of apple MADS TFs during dormancy, as previous genetic and molecular evidence highlighted their roles during the dormancy cycle [4,5,16,25,46,47]. To gain insights whether the observed physical interactions between apple MADS TFs have biological relevance during dormancy, the transcript levels of their encoding genes were quantified during an annual time course (Figure 2D). Apple buds from field-grown ‘Gala’ trees were harvested from endodormancy establishment to ecodormancy release. The accumulation of chilling hours was daily recorded in the field (Supplemental Figure S6), and a forcing test (see Methods) was performed to determine the transition from endo- to ecodormancy (mid to late January). The expression peak of *MdDAM1* occurred in the middle of endodormancy, with decreasing levels after the transition to ecodormancy. *MdDAM2* and *MdDAMb* were more expressed before endodormancy, with low levels after endodormancy establishment. *MdDAM4* and *MdFLC* presented their highest transcript levels during the transition from endo- to ecodormancy. *MdSVPa* and *MdSVPb* showed low transcriptional variations during the dormancy cycle. The expression profiles and the protein interaction network obtained for these MADS genes support the hypothesis that MdSVPa is a central hub for signaling, whereas MdDAM1, MdDAM4, MdDAMb and MdFLC act in concert with MdSVPa during specific dormancy cycle phases.

#### Transcriptional complexes containing MdSVPa bind to hundreds of apple genes

To understand how complexes containing MdSVPa regulate genome expression, seq-DAP-seq [43,55] was carried out to profile their DNA-binding sites. Full-length sequences of MdDAM1, MdDAM4, MdFLC and MdSVPa were tagged with C-terminal 3xFLAG or 5xMyC, aiming to avoid epitope tag interference to the N-terminal MADS DNA-binding domain. MdDAM2 and MdDAMb were not included in this experiment as they showed low expression levels after endodormancy establishment (Figure 2D). Protein complexes were co-produced by coupled *in vitro* transcription-translation, purified using anti-FLAG beads, and visualized in Western blots after coimmunoprecipitation. The interactions previously obtained were validated besides the identification of homodimer formation for MdDAM1, MdDAM4, MdFLC and MdSVPa (Supplemental Figure S7A). Next, MdDAM1-, MdDAM4, or MdFLC-MyC were independently co-produced *in vitro* with MdSVPa-FLAG, and each complex was sequentially purified using anti-FLAG and anti-MyC beads. For the homomeric complexes, only MyC-tagged protein versions were produced followed by immunoprecipitation using anti-MyC beads. Strikingly, only complexes containing MdSVPa were able to bind to a DNA probe carrying two CArG boxes, as shown by EMSA experiments (Figure 3A). In the light of these results, only protein complexes containing MdSVPa were further investigated in the seq-DAP-seq assays. MdDAM1–MdSVPa, MdDAM4–MdSVPa, MdFLC–MdSVPa and MdSVPa–MdSVPa TF complexes were produced and purified as described above, and incubated with an apple genomic DNA library obtained during dormancy. DNA fragments were recovered, sequenced using massive parallel sequencing, and aligned to the apple genome [56].

**Figure 3.**
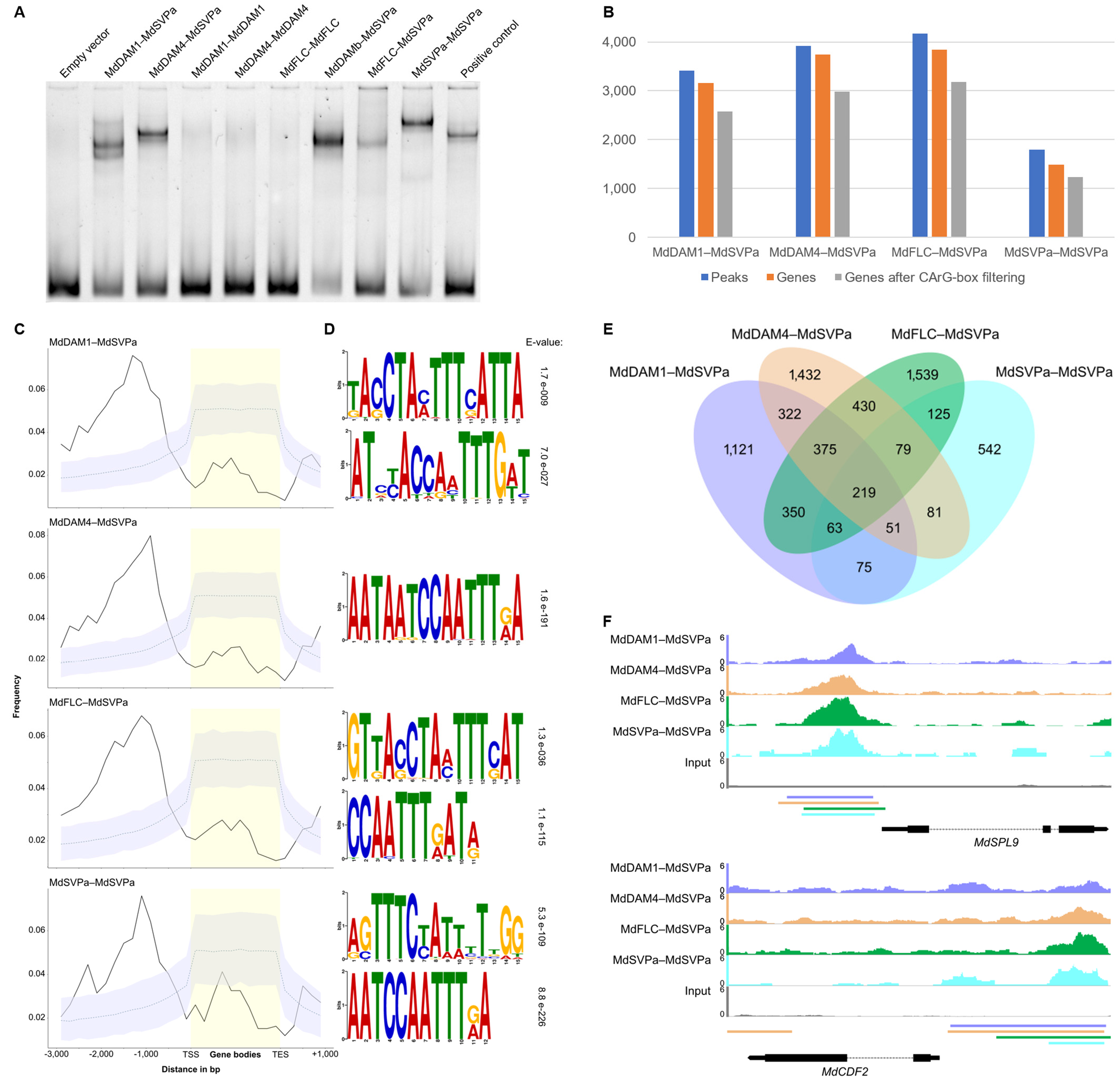
Genome-wide target sites of transcriptional complexes containing MdSVPa. **A)** EMSA assay evaluating the ability of homo- and heterodimers to recognize a DNA probe consisted of 103 bp fragment of the Arabidopsis *SEP3* promoter containing two CArG boxes [57]. The Arabidopsis AG-SEP3 complex was used as a positive control and the obtained band represents the size of an heterotetramer. **B)** Number of seq-DAP-seq peaks, peaks associated to genes and number of genes obtained after filtering for the presence of at least one CArG-box. **C)** Peak distribution in a region from 3 kb upstream of the transcription start site (TSS) to 1 kb downstream of the transcription end site (TES) of the closest gene. Solid lines represent the positional distribution of observed peaks for each complex. Dashed lines correspond to the observed positional distribution obtained from 1,000 randomly generated peak sets. As the distributions were determined using bins, the 95% confidence interval (gray shaded color) for each bin is depicted. **D)** Logos of the enriched sequence motifs identified by MEME motif analysis. The Evalue indicates the statistical significance for the corresponding motif. **E)** Venn diagram illustrating common genes between the four seq-DAP-seq datasets after filtering for the presence of CArG boxes. **F)** DNA-binding profiles of the four complexes and the control (input) to the promoter region of *MdSPL9* and *MdCDF2*. Horizontal bars below the plots represent the position of the peak regions. The color code between the plots and the bars is preserved. The Integrated Genome Browser (IGB) was used for visualization.

For MdDAM1–MdSVPa, MdDAM4–MdSVPa and MdFLC–MdSVPa, 3,407, 3,922 and 4,167 merged peaks were called, which were assigned to 3,161, 3,738 and 3,841 neighboring genes, respectively (Figure 3B; Supplemental Data S1). Fewer peaks (1,801) and gene models (1,485) were identified for the MdSVPa homodimer. Genomic enrichment upstream of the transcription start site (TSS) was observed for all transcriptional complexes, consistent with a function as TFs (Figure 3C). Two independent strategies were followed to identify the most enriched cis-elements bound by each transcriptional complex. *De novo* motif discovery detected an enrichment of two putative CArG boxes with similar nucleotide composition but different number of consecutive nucleotides in the central A/T-rich region (Figure 3D). Sequences similar to the canonical CArG-box (CC(A/T)6GG) were observed for MdDAM1–MdSVPa, MdFLC–MdSVPa and MdSVPa–MdSVPa complexes, whereas a motif containing 5 bp-length in its A/T-rich region (CC(A/T)5GG) was recurrently detected in all complexes using several independent motif discovery algorithms. CArG-box with variable A/T-length content were previously reported as DNA-binding sites of MADS TFs [42,52], including SVP and FLC [36]. Alternatively, a manual search for CArG-box motifs was performed (see Methods), and 80% of the peaks mapped close to gene models contained at least one CArG-box (Figures 3B and Supplemental Figure S7B, Supplemental Data S2).

A comparison between the shared target genes of the four complexes after the CArG-box filtering revealed that 55% of the targets were common to at least two complexes (Figure 3E). 219 target genes were bound by all four complexes, including important flowering-time regulators such as *SQUAMOSA PROMOTER-BINDING PROTEIN-LIKE 9 (MdSPL9*, MD14G1060200) and *CYCLING DOF FACTOR 2 (MdCDF2*, MD16G1071400; Figure 3F). Interestingly, the dormancy-related genes *MdDAM1, MdDAMb* and *MdFLC* were found as targets of MADS TF complexes. *MdDAM1* was bound by MdDAM4–MdSVPa, MdFLC–MdSVPa and MdSVPa–MdSVPa complexes, *MdDAMb* was a target of MdFLC–MdSVPa, whereas *MdFLC* was bound by MdDAM1–MdSVPa and MdSVPa– MdSVPa TFs (Supplemental Figure S7C). GO (gene ontology) term analysis showed a common enrichment in all four complexes in categories related to response to chemicals and endogenous and abiotic stimuli, cellular processes, carbohydrate metabolic processes, multicellular organism development and anatomical structure development (Supplemental Figure S7D). Terms related to flower development, reproduction, and post-embryonic development were enriched in all complexes, except MdFLC–MdSVPa. MdDAM1–MdSVPa, MdDAM4–MdSVPa and MdFLC–MdSVPa targets were enriched for cell differentiation, cell communication, signal transduction, abscission, metabolic process and transport.

#### Several biological processes are differentially regulated by each transcriptional complex

A complementary strategy was designed to evaluate the transcriptional changes of direct targets of these four MADS TFs in apple. Apple calli were transformed with constructs carrying MdDAM1, MdDAM4, MdFLC and MdSVPa fused to the glucocorticoid receptor (GR). Positive transformed calli (see Methods) for each gene construct were pretreated with the translational inhibitor cycloheximide (CHX), followed by dexamethasone (DEX) or solvent (mock) treatments [57]. RNA-seq analysis for each treatment and for each construct was conducted after calli were incubated for 8 hours at room temperature (Supplemental Figure S8A, Supplemental Data S3). Differentially expressed genes (DEGs, for adj. P ≤ 0.05) were identified by comparing DEX vs mock treatments. After DEX induction, 15,180, 6,247, 8,528, and 13,991 DEGs were detected for MdDAM1-GR, MdDAM4-GR, MdFLC-GR and MdSVPa-GR, respectively (Figure 4A). Comparisons among the shared target genes of each MADS TF showed that 79-91% of the DEGs were common to at least two TFs (Supplemental Figure S8B), suggesting that they may be acting in similar pathways. MdDAM1-GR dataset presented the highest percentage of unique targets (21%), whereas MdDAM4-GR showed the lowest percentage (9%). Next, we checked for crossed regulation among these four MADS TFs (Supplemental Data S3). *MdDAM1* was repressed by MdDAM1-GR and MdSVPa-GR, whereas *MdSVPa* was slightly induced by MdDAM1-GR. *MdDAM4* and *MdFLC* showed autoregulation, being repressed and induced by their encoded TFs, respectively. A GO analysis was performed and enriched terms were classified as up- or downregulated based on the expression of the genes that composed each category (Supplemental Figure S8C, see Methods). All libraries showed repressed GO terms related to secondary metabolic processes, lipid metabolic processes, abscission, cell death, and response to stress, to chemicals, and to biotic, abiotic and external stimuli; whereas translation, embryo development, multicellular organism development and anatomical structure development were induced. Some categories such as signal transduction and cell communication showed differential regulation, being induced in MdDAM4-GR and MdFLC-GR while being repressed in MdDAM1-GR and MdSVPa-GR datasets. An opposite trend was observed for carbohydrate metabolic processes and metabolic processes. This analysis indicates that several processes are being commonly regulated by the different TFs.

**Figure 4.**
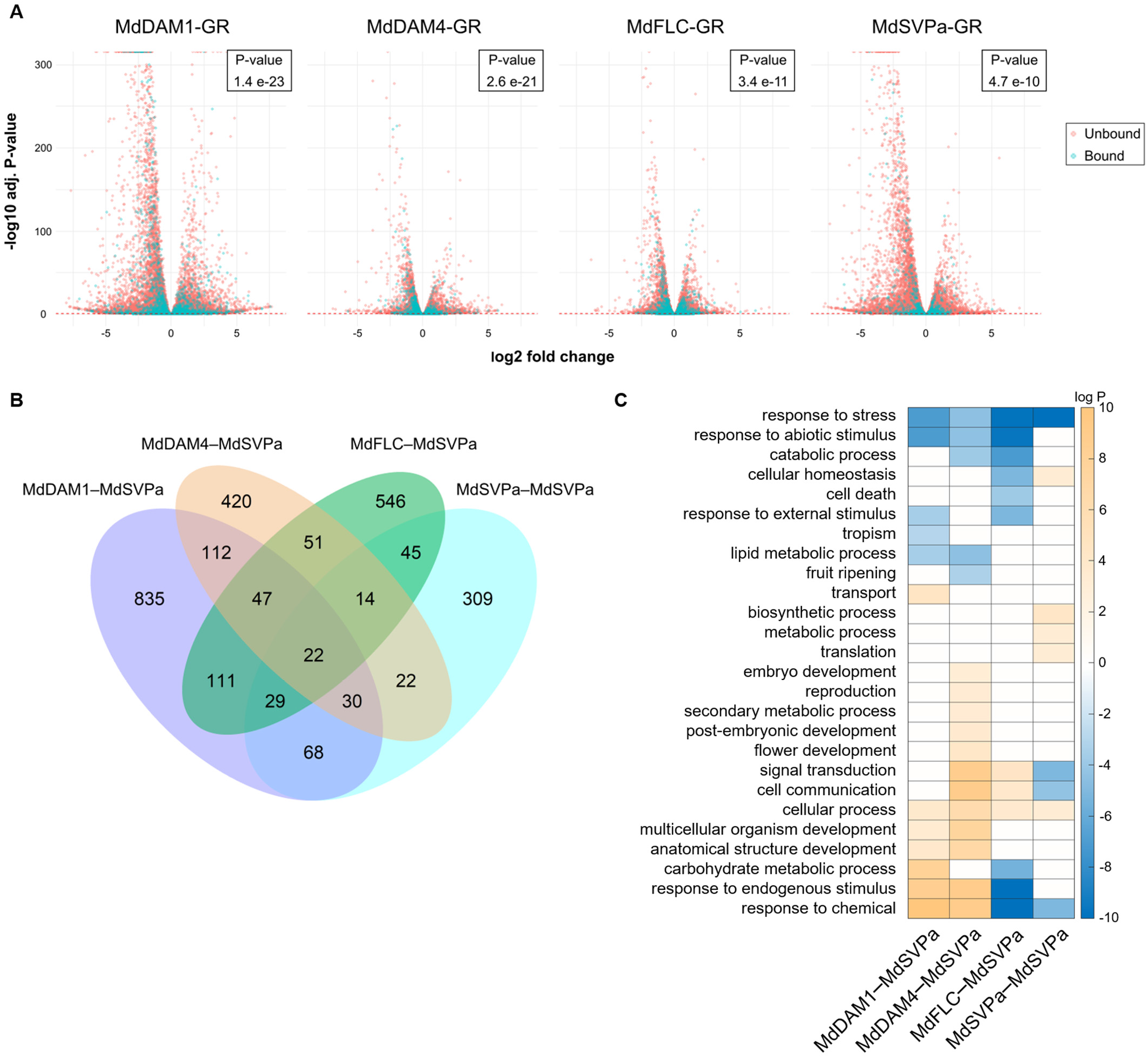
Characterization of the target genes from four transcriptional complexes containing MdSVPa. **A)** Volcano plots representing DEGs identified using the GR DEX-inducible system in transgenic apple calli transformed with four apple TFs. Bound genes represent genes also identified using the seq-DAP-seq strategy. The inset shows the P-value (one-sided Fisher’s exact test) obtained when analyzing the statistical significance of the overlap between target genes identified using seq-DAP-seq and the DEX-inducible assay. **B)** Venn diagram illustrating common genes between the high-confidence targets of each complex. **C)** GO term enrichment analysis of the high-confidence targets of each transcriptional complex containing MdSVPa. Enrichment tests were performed separately for up- and downregulated gene sets and only the best P-value (smallest) was kept. For data visualization, the best P-value was transformed using -log or log when it belonged to the up (orange gradient) or downregulated (blue gradient) gene set, respectively. Note that all P-values higher than 0.05 were replaced by zero (white boxes).

The seq-DAP-seq data obtained for the four complexes containing MdSVPa were then compared with the DEGs from the GR DEX-inducible assay. A statistically significant overlap (one-sided Fisher’s exact test, inset in Figure 4A) was obtained for all comparisons, with 54%, 28%, 35% and 51% of the genes with peaks called in the seq-DAP-seq of MdDAM1–MdSVPa, MdDAM4–MdSVPa, MdFLC–MdSVPa and MdSVPa–MdSVPa, respectively, also being differentially expressed in the GR DEX-inducible experiment. The genes found in the intersection between the two independent assays were considered as high-confidence targets of each complex (Supplemental Data S4). Further analysis of these high-confidence datasets showed that 56-70% of the DEGs with peaks were repressed in the DEX-inducible system (Figure 4A). GO term analysis provided further evidence that the different protein complexes containing MdSVPa differentially regulate plant processes (Figure 4C). All complexes repressed genes belonging to response to stress, and induced genes related to cellular processes. MdDAM1–MdSVPa and MdDAM4–MdSVPa induced target genes related to multicellular organism development, anatomical structure development, and response to endogenous stimulus and to chemicals, whereas they repressed genes related to lipid metabolic processes and response to abiotic stimulus. Additionally, MdDAM4–MdSVPa targets were enriched with induced categories related to embryo development, reproduction, secondary metabolic processes and flower development. MdFLC–MdSVPa repressed genes related to carbohydrate metabolic processes, cell death, cellular homeostasis, and response to chemicals, and to endogenous, abiotic, and external stimuli. On the other hand, the MdSVPa homodimer induced genes related to cellular homeostasis, biosynthetic and metabolic processes, and translation, whereas repressed genes associated with signal transduction, cell communication and response to chemicals.

#### Targets from complexes containing MdSVPa but not the MdSVPa homodimer are enriched with DEGs during the dormancy cycle

As the genes coding for MdDAM1, MdDAM4 and MdFLC were seasonally regulated during bud dormancy (Figure 2D), we tested whether the high-confidence targets of each complex were enriched for genes with differential expression during the dormancy cycle. For this purpose, we re-analyzed two public RNA-seq experiments performed with apple terminal buds harvested in field conditions (dataset A) [4] or under controlled temperatures (dataset B) [5]. After quantifying the expression of transcripts in both datasets (see Methods), the expression profiles obtained for *MdDAM1, MdDAM4, MdFLC* and *MdSVPa* were similar to the ones found in our experiment (Figure 2D and Supplemental Figure S9A-B), validating the further use of these datasets. For dataset A, pairwise comparisons were performed between dormant samples harvested from October to February (endo- to ecodormancy transition) in comparison to budbreak samples harvested in March. In total, 7,302 DEGs (for adj. P ≤ 0.05 and fold2change < or > 1.75) were identified in at least one of the comparisons, and were used in an enrichment analysis. Notably, the high-confidence targets of MdDAM1–MdSVPa, MdDAM4–MdSVPa, and MdFLC–MdSVPa were enriched with DEGs from dataset A (hypergeometric test, for P ≤ 0.05; Figure 5A). A similar strategy was employed for dataset B. Samples harvested 10, 25, 35 and 65 days after cold treatment were compared to endodormant samples obtained in the field before artificial cold exposure. Overall, 4,080 DEGs (for adj. P ≤ 0.05 and fold2change < or > 1.75) were identified in at least one of the pairwise comparisons. As observed in the previous analysis, only targets from MdDAM1–MdSVPa, MdDAM4–MdSVPa, and MdFLC–MdSVPa complexes were enriched with DEGs from the analysis of dataset B (hypergeometric test, for P ≤ 0.05; Figure 5B). Conversely, target genes from the MdSVPa homodimer were not enriched with DEGs during dormancy in any dataset (Figure 5A-B), which suggests that the role of MdSVPa as regulator of dormancy requires its association with other MADS TFs.

**Figure 5.**
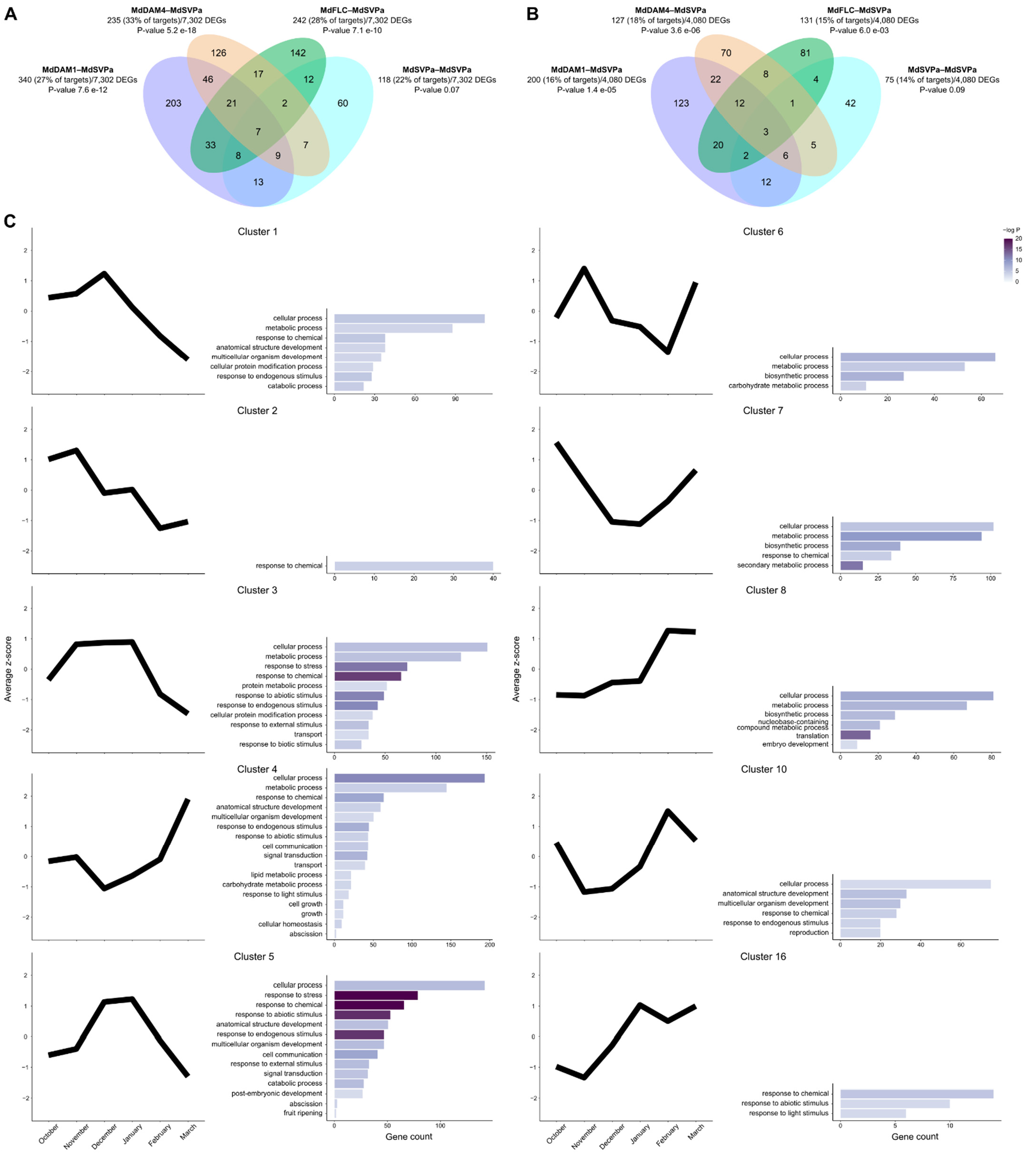
Comparisons between dormancy-related DEGs and the target genes of four transcriptional complexes containing MdSVPa. **A)** Venn diagram showing the overlap between dormancy-related DEGs identified in field-grown samples and the high-confidence target genes of each transcriptional complex. A gene was considered as differentially regulated during dormancy (adj. P ≤ 0.05 and fold2change < or > 1.75) if present in at least one time point in comparison to budbreak [4]. The target gene datasets of each transcriptional complex were evaluated for the enrichment of dormancy-related DEGs, and the obtained P-value (hypergeometric test) is shown. **B)** Venn diagram displaying the overlap between dormancy-related DEGs identified under artificial cold exposure and the high-confidence target genes of each transcriptional complex. A gene was considered as differentially expressed (adj. P ≤ 0.05 and fold2change < or > 1.75) if present in at least one time point in comparison to endodormancy establishment [5]. The lists of target genes from each transcriptional complex were evaluated for the enrichment of dormancy-related DEGs, and the obtained P-value (hypergeometric test) is shown. **C)** Gene expression and GO enrichment analysis of the high-confidence target genes of MdDAM1, MdDAM4 and MdFLC in association with MdSVPa during dormancy (Supplemental Data S5). Average z-score values for clusters were obtained from an annual time course expression analysis of apple buds harvested from field-grown ‘Golden delicious’ trees according to Moser et al [4]. Only clusters showing at least one significant GO term (P-value < 0.05) are represented.

To further probe the role of heteromeric complexes during dormancy, we generated a list containing all target genes of MdDAM1, MdDAM4 and MdFLC in transcriptional complexes with MdSVPa (2,356 unique genes) and analyzed their expression dynamics during a dormancy cycle (2,324 out of 2,356 genes were present in dataset A, [4]). Based on their expression patterns, 17 clusters were identified (Supplemental Data S5). Complementarily, a GO term enrichment analysis was performed to explore the pathways and functions associated with each cluster (Figure 5C). Genes from cluster 4, 7 and 10 were repressed during endodormancy, and GO categories related to response to chemicals and cellular processes were common to all three clusters. Cluster 4 was the only one showing terms associated with growth, cell growth and cellular homeostasis, whereas terms associated with response to endogenous and abiotic stimuli, abscission, signal transduction and cell communication were also enriched in cluster 5, and clusters 1 and 3 to some extent. Interestingly, these three clusters showed the exact opposite expression pattern of cluster 4 during dormancy, i.e. a peak of expression during endodormancy. Moreover, cluster 5 uniquely presented terms related to post-embryonic development and fruit ripening. Taken together, these analyses strongly suggest that MdSVPa, when associated with other MADS TFs, has a regulatory role in several plant processes that occur during the dormancy cycle. However, MdDAM1– MdSVPa, MdDAM4–MdSVPa, and MdFLC–MdSVPa transcriptional complexes showed a high number of unique dormancy-related DEGs (Figure 5A-B), suggesting that each complex regulates different processes of the dormancy cycle.

#### Genes showing similar expression profile but regulated by different MdSVPa-containing complexes have distinct biological functions during dormancy

A similar approach was employed for the individual target genes of MdDAM1–MdSVPa, MdDAM4– MdSVPa, and MdFLC–MdSVPa complexes aiming to identify the individual contribution of each complex to dormancy control. Based on the expression patterns of MdDAM1–MdSVPa targets during dormancy, 11 clusters were identified (Supplemental Data S6) and five of them were enriched in a GO term analysis (Figure 6A). Cluster 2 and 3 shared GO terms associated with cellular processes, regulation of molecular function, anatomical structure development, and response to chemicals and endogenous stimulus. However, these clusters presented antagonistic patterns of expression, as cluster 2 showed high expression levels during endodormancy, while cluster 3 was repressed at the same time points. Additionally, cluster 2 was enriched with terms related to circadian rhythm, post-embryonic development, and response to abiotic and light stimuli. Conversely, cluster 3 showed terms associated with growth, cell communication, and signal transduction. Cluster analysis for the target genes of MdDAM4–MdSVPa identified nine clusters during the dormancy cycle (Figure 6B and Supplemental Data S7). Genes belonging to cluster 1 were induced near budbreak, with low levels during endodormancy. Moreover, this cluster was enriched with terms associated with response to chemicals and endogenous stimulus, signal transduction, cell communication, and flower development. Interestingly, cluster 4 was enriched with the same GO terms, except flower development, besides response to stress and abiotic stimulus. The genes that composed cluster 4 showed a seasonal accumulation of transcripts during winter, in an opposite trend to the one observed to cluster 1. Finally, the high-confidence targets of MdFLC–MdSVPa were classified in 12 clusters based on their expression dynamics during dormancy (Supplemental Data S8), and seven of them were enriched after a GO analysis (Figure 6C). Genes composing cluster 1 were upregulated during endodormancy, and categories associated with circadian rhythm, signal transduction, cell communication, and response to external, endogenous and abiotic stimuli were found enriched.

**Figure 6.**
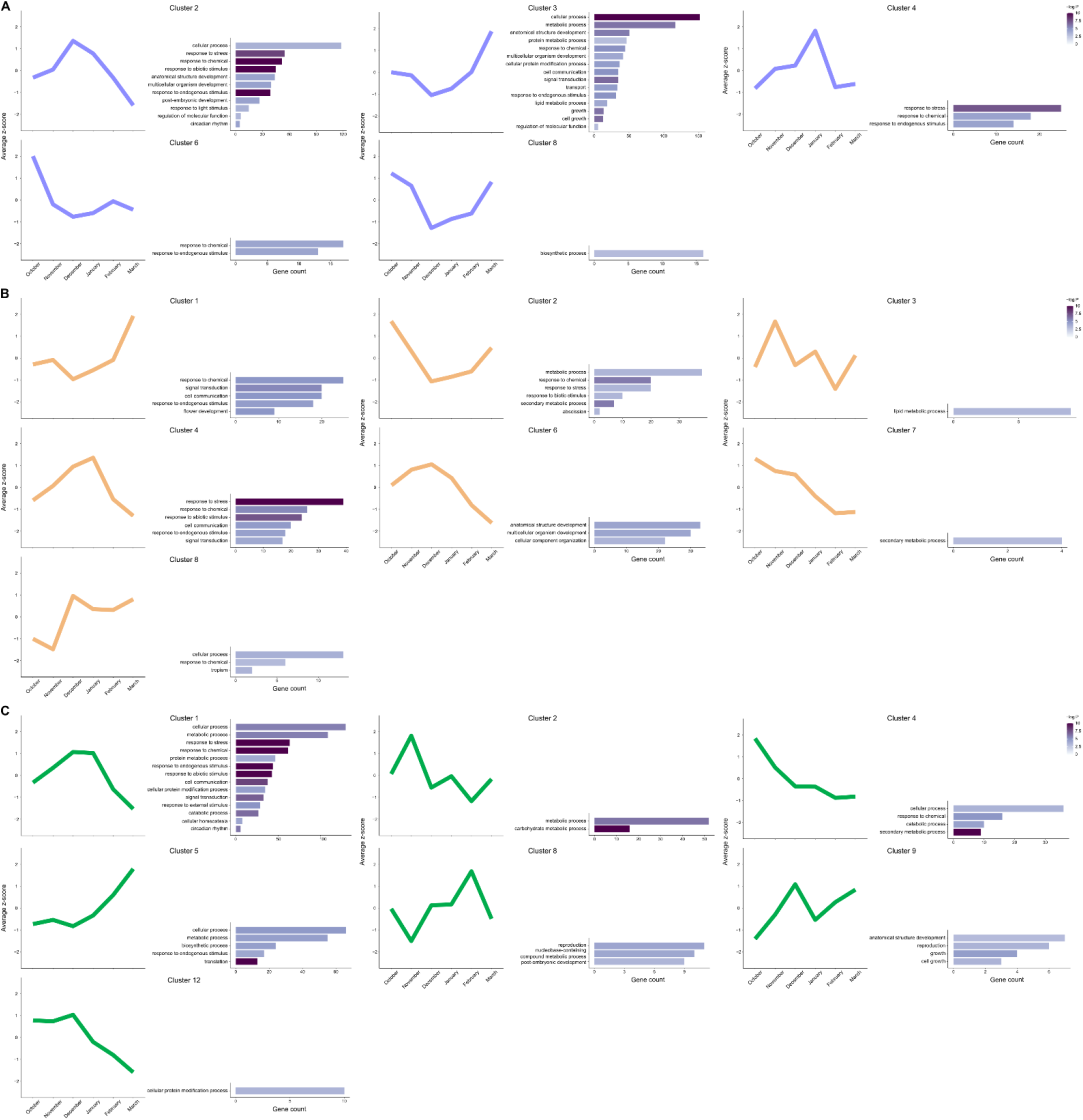
Average expression patterns and GO enrichment analysis of the different clusters obtained during apple bud dormancy. **A)** Analysis of the high-confidence target genes of MdDAM1–MdSVPa during dormancy (Supplemental Data S6). **B)** Analysis of the high-confidence target genes of MdDAM4–MdSVPa during dormancy (Supplemental Data S7). **C)** Analysis of the high-confidence target genes of MdFLC–MdSVPa during dormancy (Supplemental Data S8). Average z-score values for clusters were obtained from an annual time course expression analysis of apple buds harvested from field-grown ‘Golden delicious’ trees according to Moser et al [4]. Only clusters showing at least one significant GO term (P-value < 0.05) are represented.

The analyses performed so far focused on identifying general GO categories, as we used a subset of GO terms called ‘plant GO slim’ for filtering the enrichment assays. However, even though clusters from different complexes showing a similar expression profile also shared similar GO terms, specific biological functions could be assigned to each cluster. One example is cluster 2 from MdDAM1–MdSVPa and cluster 4 from MdDAM4–MdSVPa (Figure 6A-B), which showed a peak of transcripts during endodormancy and low expression levels close to budbreak. Both clusters were enriched for categories associated with response to stress, to chemicals, and to abiotic and endogenous stimuli. However, only MdDAM1–MdSVPa targets were enriched with terms related to response to light, regulation of molecular function and circadian rhythm. Conversely, targets of MdDAM4–MdSVPa showed cell communication and signal transduction as significant categories. These results suggest that each transcriptional complex is able to co-regulate genes in a similar manner, but these clusters are composed by different sets of genes and so have different biological functions during dormancy. To go deeper into the individual contributions of each transcriptional complex to dormancy control, we performed an independent enrichment analysis that only considered the broad GO domain of biological processes (Supplemental Data S9). In this analysis, four clusters of MdDAM1–MdSVPa targets were enriched. The same amount of clusters were also found enriched for the targets of MdDAM4-MdSVPa, and the target genes of the MdFLC-MdSVPa complex showed eight clusters with significant categories. Using the same example as above, cluster 2 from MdDAM1–MdSVPa and cluster 4 from MdDAM4–MdSVPa were both enriched with GO terms linked to basic cellular and metabolic processes, and regulation of transcription. However, cluster 2 showed categories related to photoperiodism, flowering, circadian rhythm, and hormonal responses (i.e. abscisic acid [ABA], jasmonate and salicylic acid stimuli), whereas cluster 4 presented terms associated with ethylene stimulus, drought and trehalose biosynthetic process.

#### MADS TF complexes act in concert to fine-tune the dormancy cycle in apple

Next, we focused on identifying genes playing key roles during the dormancy cycle and acting downstream of the investigated MADS transcriptional complexes. A significant overlap was identified between the lists of DEGs from the transcriptomic analyses done from datasets A and B (hypergeometric test, for P ≤ 0.05; Supplemental Figure S9C), and a list of DEGs found in the intersection between the two analyses was generated (dormancy-related DEGs, Supplemental Data S10). After, we obtained the overlap between the dormancy-related DEGs and the high-confidence targets of MdDAM1–MdSVPa, MdDAM4–MdSVPa, and MdFLC–MdSVPa complexes. In total, 231 genes fit this selection criteria (Supplemental Data S11), and some of these genes were further characterized based on their putative role in dormancy regulation. *MdBRC1 (BRANCHED 1*; MD06G1211100) and *MdMP (MONOPTEROS/AUXIN RESPONSE FACTOR 5;* MD15G1014400) were both bound and induced by MdDAM1– MdSVPa and MdDAM4–MdSVPa complexes (Supplemental Figure S10A). These two genes are known to be part of GRNs regulating bud and meristem development [26,58–61]. The peak of expression of *MdBRC1* occurred during the transition from endo- to ecodormancy, maintaining higher levels until budbreak (Figure 5C, cluster 16), whereas *MdMP* only showed changes in transcript levels near budbreak (Figure 5C, cluster 8). In spite of both genes being common targets of the same transcriptional complexes, they showed a differential temporal regulation of expression during dormancy. This trend highlights the complex gene regulation happening during the dormancy cycle.

Several genes related to hormone biosynthesis and signaling were also present in this list. *MdCKX5* (cytokinin oxidase/dehydrogenase 5; MD15G1021500), which transcripts accumulated during endodormancy (Figure 5C, cluster 3), was bound and induced by MdDAM1–MdSVPa and MdFLC–MdSVPa complexes (Supplemental Figure S10A). Arabidopsis CKX5, together with CKX3, regulates the activity of the reproductive meristems by catalyzing the oxidation of cytokinins (CK) [62]. Recently, it was proposed that CK stimuli repress *MdDAM1* expression [63], suggesting that a negative feedback loop between MADS TFs and CK may exist. Another gene present in cluster 3 (Figure 5C) was *MdGA2ox1 (GA2-oxidase 1*; MD05G1207000), which was bound and repressed by the MdFLC–MdSVPa TF complex (Supplemental Figure S10A). GA2ox are enzymes responsible for the deactivation of bioactive GA, preventing the accumulation of this growth-promoter hormone during dormancy [64–66]. GA function is antagonized by ABA, a hormone that promotes shoot growth cessation and bud dormancy establishment [64,65]. *MdNCED4 (9-cis-epoxycarotenoid dioxygenase 4;* MD16G1090700) and *MdPYL4 (pyrabactin resistance-like 4;* MD07G1227100) are involved in ABA biosynthesis and signaling, respectively, and showed opposite expression trends during dormancy. *MdNCED4* was repressed after dormancy establishment (Figure 5C, cluster 4), while *MdPYL4* was induced during endodormancy (Figure 5C, cluster 3). Likewise, *MdNCED4* was repressed by MdDAM1–MdSVPa and *MdPYL4* was induced by MdFLC–MdSVPa (Supplemental Figure S10A). Similar expression profiles during dormancy were described for these classes of genes in pear [67].

We have also observed the presence of genes related to cell wall modification processes. The involvement of xyloglucan metabolism during the dormancy cycle was already reported elsewhere [68–70]. *MdXTH9 (XTH* for *xyloglucan endotransglucosylase/hydrolase;* MD09G1102600), *MdXTH15* (MD09G1152700) and *MdXTH23* (MD13G1237300), three genes encoding enzymes related to loosening and rearrangement of the cell wall, were induced by MdDAM1–MdSVPa complexes (Supplemental Figure S10A). Conversely, *MdXTH15* was repressed by the other MADS complexes. Interestingly, poplar plants ectopically expressing the peach TF *EBB1* (*EARLY BUD BREAK 1*) showed early budbreak, and several genes related to cell wall modifications such as *XTH9* were upregulated in the transgenic plants in relation to WT ones [71].

Finally, some genes related to flowering-time regulation through the photoperiodic pathway also fit our selection criteria [72,73]. *MdCDF2* and *MdLHY (LATE ELONGATED HYPOCOTYL*, MD01G1090900) were upregulated during endodormancy, showing lower expression levels near budbreak (Figure 5C, cluster 3). *MdCDF2* was bound and slightly repressed by MdDAM4–MdSVPa, whereas *MdLHY* was induced by MdDAM1– MdSVPa and repressed by MdDAM4–MdSVPa (Figure 3F and Supplemental Figure S10A). In Arabidopsis, both LHY and CDFs act in overlapping flowering pathways, leading to the activation and accumulation of CONSTANS (CO), which further activates the expression of *FT* and, consequently, *SOC1* [74,75]. Remarkably, *MdSOC1a* (MD02G1197400), an ortholog of the Arabidopsis floral integrator gene *SOC1* [29], was bound and repressed by the MdSVPa homomeric complex (Supplemental Figure S10B). In Arabidopsis, SVP represses *SOC1* expression as a part of a GRN that regulates floral transition in the SAM [29]. The SVP-SOC1 regulatory module seems conserved in apple, although its implication in flowering and/or dormancy control remains unknown.

### Discussion

The finely tuned control of dormancy is fundamental for temperate fruit trees to ensure their survival during winter, while being ready to maximize their reproductive cycle after budbreak. Recent genetic studies revealed that DAM- and SVP-like MADS TFs act as master regulators of the dormancy cycle in Rosaceous fruit trees. However, how they integrate molecular networks to shape dormancy dynamics has not been properly assessed yet. Here, we addressed this question by employing for the first time in non-model species seq-DAP-seq, a tool that allows the identification of target genes specific to transcriptional complexes. Moreover, by coupling it to GR DEX-inducible assays, we managed to partially overcome the difficulties associated with doing functional genetics in fruit tree species.

#### Apple MADS TF complexes operate during the dormancy cycle of fruit trees

We have rationalized that MADS TFs involved in dormancy cycle in trees form molecular complexes whose function is defined by their combinatory composition. Consequently, we have demonstrated the existence of apple MADS TF complexes by different means, i.e. Y2H and *in vitro/in planta* co-IP assays, and found that MdSVPa is a central component of these complexes (Figures 2A-C and Supplemental Figure S7A). Indeed, the role of MdSVPa seems crucial for the transcriptional function of these multimeric complexes, as the absence of MdSVPa compromises their ability to bind to DNA (Figure 3A). The relative abundance of these complexes during the dormancy cycle might depend on the expression levels of their encoding genes. In Arabidopsis, SVP forms complexes with other MADS TFs, such as FLC, FLM, AGL24, FUL and SOC1, to regulate vegetative development, reproductive transition and flower development [51]. This functional plasticity is partly conferred by the temporal and spatial pattern of expression of *SVP* mRNA. *SVP* is expressed in the vegetative meristem where its protein product interacts with FLC to repress floral transition [29]. After floral transition, *SVP* is expressed in the floral meristem where SVP forms complexes with AP1 and AGL24 to regulate flower development [76,77]. ChIP-seq and transcriptomic experiments showed that switching molecular partners results in distinct SVP DNA-binding specificity, leading to the expression and/or repression of different sets of target genes [27,36]. Our results indicate that *MdSVPa* is expressed at constant levels in apple winter buds, which is compatible with MdSVPa TF being part of molecular complexes along all the dormancy cycle progression (Figure 2D and Supplemental Figure 9A-B). Remarkably, *MdDAM1* and *MdDAM4* showed their maximal gene expression in the endodormant phase, whereas *MdFLC* mRNA peaked at the transition from endo- to ecodormancy (Figure 2D). Therefore, we propose a model in which MdSVPa sequentially forms complexes with the MADS TFs that predominate at each dormancy-cycle phase (Figure 7). The composition of these complexes could alter their DNA-binding specificity and, therefore, the transcriptional regulation of their target genes.

**Figure 7.**
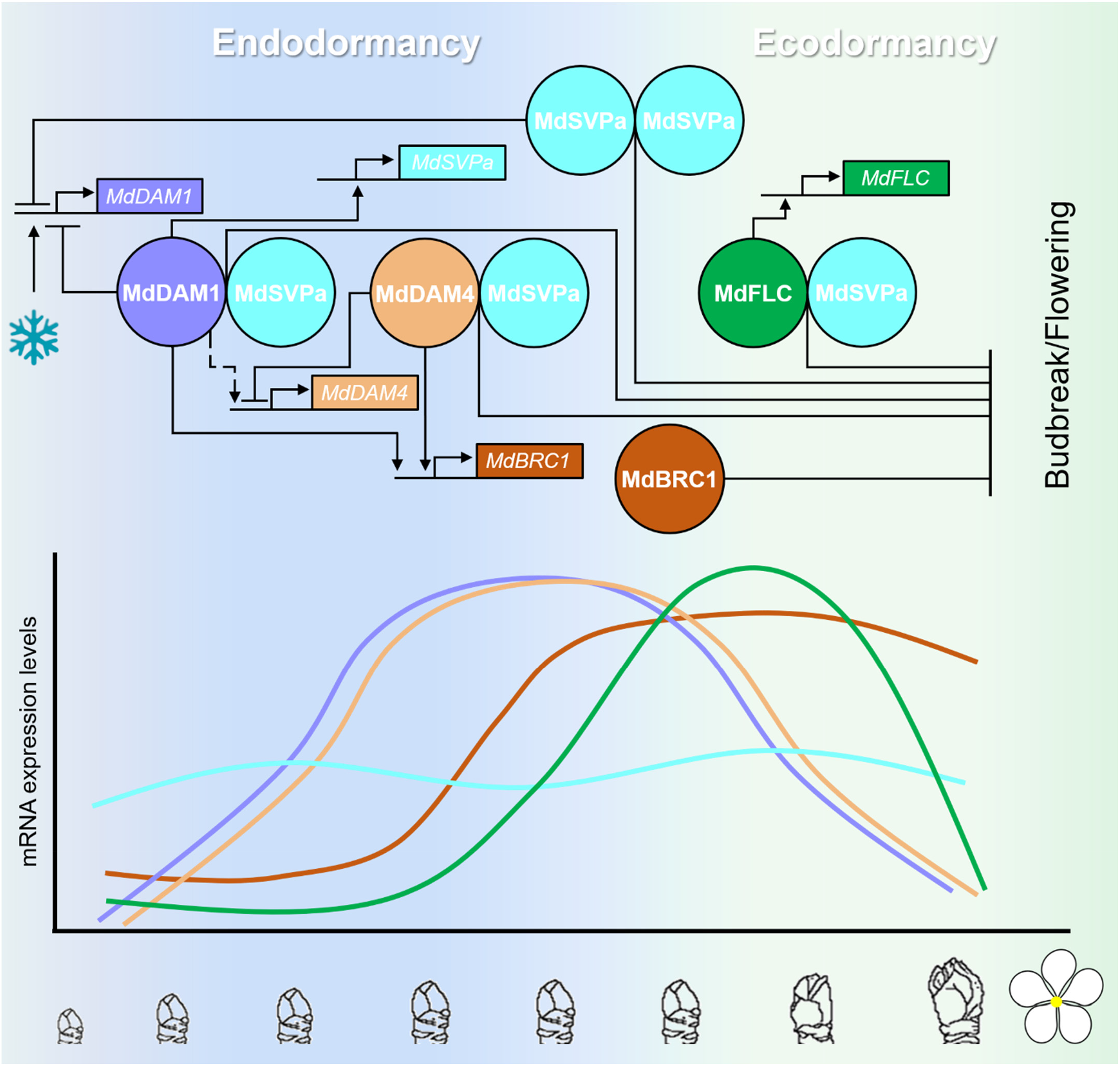
Tentative model summarizing the regulatory interactions between MADS TFs during the dormancy cycle. Apple MADS TFs MdDAM1, MdDAM4, MdFLC and MdSVPa form complexes and regulate the expression of each other in a genetic regulatory circuit that integrates environmental signals to restrict bud growth during winter dormancy. We propose that this GRN leads to budbreak and flowering inhibition at least partially through the transcriptional control of *MdBRC1*. Arrowed lines indicate transcriptional activation whereas blunted lines mean transcriptional repression. The dotted line represents the induction of *MdDAM4* by MdDAM1 as proposed by [4]. Genes and proteins are represented by boxes and circles, respectively. In the lower part of the cartoon, the mRNA expression level for each gene is represented (color code as shown in the upper part).

To address this question, we have exploited the recently developed technology called seq-DAP-seq [43]. Compared to other approaches, such as ChIP-seq and DAP-seq, seq-DAP-seq allows the identification of genome-wide binding sites of multimeric complexes. As seq-DAP-seq does not require either creating transgenic lines or generating antibodies, it can be easily applied to any plant species. However, seq-DAP-seq has been only used to study TFs in the model plant species Arabidopsis [43]. Here, we have demonstrated that seq-DAP-seq is a useful approach to determine genome-wide binding sites of multimeric MADS TFs in fruit trees. As expected, apple MADS complexes preferably bind to DNA regions containing CArG-boxes (Figure 3C and Supplemental Figure S7B). A detailed analysis showed that MADS TF combinatory dimerization results in a preference for specific CArG-box displaying slightly altered nucleotide composition and A/T core length (Figure 3C). This altered binding preference could explain differences in target selectivity. Indeed, although many target genes were shared between the studied apple MADS complexes, a significant number of target genes were bound by a specific complex (Figure 3E-F and Supplemental Figure S10). Furthermore, by making use of the GR system [57,78], we were able to isolate genes that are bound and transcriptionally regulated by the different MADS complexes (Figure 4). The GO categorization of these target genes evidences the involvement of the studied apple MADS TFs in several biological processes during the dormancy cycle. Some of these processes are affected by a single MADS TF complex, whereas others are influenced by many of them. These results enable us to conclude that MADS complexes containing MdSVPa are sequentially formed during dormancy to modulate transcriptional responses specific to each dormancy phase (Figure 7).

#### A genetic network governed by SVP-containing complexes controls the dormancy cycle

The target genes of the studied MADS complexes might be part of GRNs regulating the dormancy cycle in apple trees. We have studied this question by making use of published RNA-seq data on apple buds during dormancy [4,5]. Remarkably, we have found a statistically significant overlap between the DEGs during dormancy (from the two unrelated RNA-seq experiments) and the high-confidence target genes of MdDAM1–MdSVPa, MdDAM4–MdSVPa and MdFLC–MdSVPa multimeric complexes (Figure 5A-B). This result indicates a relevant function of these MADS complexes in regulating dormancy in apple. Nevertheless, target genes of MdSVPa– MdSVPa homomeric complex were not statistically enriched for dormancy-related DEGs (Figures 5A-B), suggesting that MdSVPa do not have a main role as dormancy-cycle regulator and/or this role requires its association with other MADS TFs. Supporting this idea, the overexpression of *MdSVPa* delays budbreak but does not affect dormancy entrance in apple [25], probably because of the low expression levels of other MADS genes (e.g. *MdDAM1, MdDAM4* and *MdFLC*) during the autumn. Moreover, apple trees in which *MdDAM1* was silenced lost their capacity to enter into dormancy [4], although *MdSVPa* mRNA is known to be expressed all over the dormancy cycle in natural conditions (Figure 2D and Supplemental Figure 9A-B, [24,25]). Within this context, MdSVPa would be required by these MADS multimeric complexes for the proper regulation of dormancy-related genes. When analyzing the expression dynamics of the target genes from heteromeric MdSVPa-containing TF complexes, we observed the formation of gene clusters especially linked to endo- and ecodormancy (Figure 5C). Two contrasting expression patterns were recurrently identified, i.e. a peak of expression during endodormancy (clusters 1, 3 and 5), and low transcript levels during winter followed by induction of gene expression in ecodormancy and/or close to budbreak (clusters 4, 7 and 10). A recent study encompassing the transcriptomic changes happening during dormancy in sweet cherry identified similar clusters, besides additional ones related to other bud developmental stages [3]. The authors proposed a list of TFs and promoter motifs overrepresented in specific clusters that could be used as markers of some dormancy-specific phases. Some of these markers were also found within the gene clusters shown in Figure 5. Cluster 1 and 3 contained *MdABF2* (MD15G1081800) and *MdOBP1* (MD08G1040100) (Supplemental Data S4), respectively, which were proposed as markers for clusters with maximal expression during endodormancy [3]. Additionally, cluster 7 (Figure 5C) showed a gene marker (*MdDAMb*) related to maximum expression levels during paradormancy and ecodormancy. These findings further corroborate that MdSVPa-containing complexes are acting in the regulation of particular processes during the dormancy cycle, especially endo- to ecodormancy transition.

In several temperate tree species, *DAM* genes are transcriptionally induced in response to cold by the dehydration-responsive element binding (DREB) protein/C-repeat binding factors (CBFs) [79,80]. Therefore, the cold-induced transcriptional activation of *MdDAM1* and *MdDAM4* (Figure 2D) might be a prerequisite for the formation of MADS complexes with MdSVPa to activate and maintain endodormancy. In such a scenario, apple MADS complexes could operate as key regulators and integrators of environmental responses (Figure 7). Once CR is satisfied, and *MdDAM1* and *MdDAM4* mRNA expression is low, MdDAM1–MdSVPa and MdDAM4–MdSVPa complexes would be less abundant leading to endodormancy release and the subsequently transition to ecodormancy. Then, the MdFLC–MdSVPa complex would predominate until *MdFLC* gene expression is repressed at the springtime. As an additional layer of regulation, relative abundance of MdSVPa-containing complexes might be post-translationally stabilized or destabilized by low or warm temperatures, respectively, as shown for SVP complexes in Arabidopsis [38]. Therefore, apple MADS complexes are potential integrators of temperature signals into GRNs.

To elucidate these GRNs, we have individually classified the target genes of MdDAM1–MdSVPa, MdDAM4–MdSVPa and MdFLC–MdSVPa complexes in co-expression clusters during dormancy and analyzed their GO term enrichment (Figure 6). Interestingly, several clusters are enriched in categories related to development. These clusters were distributed along the target genes of the three MADS complexes, reflecting a possible reduction of meristematic activity during endodormancy followed by its reactivation in the ecodormancy and at the initiation of budbreak and/or flowering. Remarkably, the expression profile of MdDAM1–MdSVPa and MdDAM4–MdSVPa target genes related to development trend to have either their maximum or their minimum expression levels at the end of endodormancy (i.e. December). This is the precise moment at which the mRNA expression levels of *MdDAM1* and *MdDAM4* peak before their rapid downregulation (Supplemental Figure S9A). Notably, MdFLC–MdSVPa target genes related to development (Figure 6C, cluster 8) peak during ecodormancy (i.e. February), following a similar pattern of expression to that shown by *MdFLC* (Supplemental Figure S9A). These results support the notion that MADS complexes directly regulate the transcription of genes related to tree developmental processes during winter dormancy cycle. We were also interested in processes related to growth, as the endodormant phase is characterized by a reduction of bud growth activity [11]. The GO analysis showed that MdDAM1–MdSVPa complex inhibits the expression of a set of genes related to “growth” and “cell growth” at the moment of highest level of *MdDAM1* mRNA expression (Figure 6A, cluster 3). This suggests a key role of MdDAM1–MdSVPa complex in growth repression during endodormancy, which is supported by the evergrowing phenotype displayed by transgenic apple trees silencing *MdDAM1* mRNA expression [4]. Besides growth and development, many other biological processes related to the dormancy cycle were enriched in the clusters summarized in Figure 6. Numerous GO categories associated with response to stress and hormones (ABA, CK, among others), cell wall modifications, carbohydrate metabolism, and signaling were identified (Supplemental Data S9), which association to the dormancy cycle has been reported elsewhere [65].

A list of 231 MADS complexes’ target genes with potential function in the dormancy cycle was produced (Supplemental Data S11). This list contains genes whose homologues of Arabidopsis play roles in bud and meristem development (e.g. *BRC1* and *MP*), hormone homeostasis and signaling (e.g. *CKX5*, *GA2ox1*, *NCED4* and *PYL4*), cell wall remodeling (e.g. *XTH9, XTH15* and *XTH23*) and flowering-time control (e.g. *SOC1*, *LHY* and *CDF2*). Notably, homologues of the Arabidopsis gene *BRC1* are known to act as bud outgrowth repressors in several plant species [58,81]. A remarkable case is *BRC1* of hybrid aspen, in which Singh et al. [26] characterized a genetic network that mediates the control of budbreak. In this network, SVL (a SVP-like TF) is positioned upstream of *BRC1* and the GA and ABA signaling cascades. The direct binding of SVL to the *BRC1* locus induces its transcriptional activation, and *BRC1* overexpression leads to late budbreak. Therefore, the module SVL-BRC1 is believed to operate in buds to block outgrowth. In turn, the expression of *SVL* and *BRC1* is downregulated by cold, explaining, at least in part, a mechanism of temperature-mediated regulation of budbreak in trees [26]. In agreement with the mechanism described for hybrid aspen, we found that MdDAM1–MdSVPa and MdDAM4– MdSVPa complexes bind to *MdBRC1* and induce its mRNA expression (Supplemental Figure S10A). Furthermore, the expression level of *MdBRC1* (Figure 5C, cluster 16) is upregulated at the moment when *MdDAM1* and *MdDAM4* display their highest gene levels of expression (Supplemental Figure S9A). Similar gene expression dynamics during dormancy was previously reported for *MdBRC1* [23]. However, this dynamic was not disrupted in evergrowing-like RNAi plants simultaneously targeting several apple *DAM*- and *SVP*-like genes, suggesting that other pathways can also be held accountable for *MdBRC1* transcriptional modulation. Although the MdBRC1 role has not been proved yet in apple trees, it is very likely that it acts as a budbreak repressor, as reported in most of the plant species in which it has been studied so far [81,82] (reviewed by [58]). In apple, the high levels of *BRC1* expression observed during ecodormancy (Figure 5C, cluster 16), after CR is satisfied, could indicate a main function during this dormancy phase. Strikingly, the *BRC1* homologue in the perennial plant *Arabis alpina* (*AaBRC1*) plays an important role in maintaining bud dormancy during and after vernalization [82]. Thus, we propose that MdBRC1 functions downstream of MdSVPa-containing complexes to repress budbreak during ecodormancy (probably together with MdFLC–MdSVPa complex). This function could be integrated in a GRN in which *MdBRC1* transcription is partially controlled by MdSVPa-containing complexes (Figure 7). The described GRN could be partially regulated by feedback loops between MADS complexes and the hormonal and environment-mediated transcriptional modulation of its key components.

#### *A neofunctionalization of* DAM *genes could cause their dormancy specialization in fruit trees*

DAM TFs are phylogenetically related to SVP from Arabidopsis and, therefore, are often referred to as SVP-like TFs. Eudicots’ SVP-like proteins have been classified in three different clades, probably originated from a whole genome triplication [83,84]. Apple DAM TFs, together with other DAM proteins from peach, plum and pear, belong to a Rosaceae-specific clade that is separated from the cluster containing the Arabidopsis SVP, which also includes MdSVPa and MdSVPb [11,83,84]. It has been proposed that these two groups were originated by a lineage-specific whole genome duplication event [83,84]. The divergence between DAM and SVP TFs is partially caused by structural changes in the DNA-binding MADS domain [85,86], suggesting functional diversification between these two groups. Here, we have provided several lines of molecular evidence that corroborate the DAM-SVP functional diversification. First, the misexpression of *MdDAM* and *MdSVP* genes from the Arabidopsis *SVP* promoter showed that only *MdSVPa* and *MdSVPb* are able to recapitulate the WT flowering phenotype of a *svp* null Arabidopsis mutant (Figure 1). By simultaneously co-expressing *MdDAM* and *MdSVP* genes in the *svp-41* mutant, we ruled out the possibility that the lack of flowering activity of MdDAM TFs was due to their inability to form functional complexes in the absence of *SVP* (Supplemental Figure S5). These results indicate that the apple and Arabidopsis *SVP* genes keep an ancestral flowering repressive function that the apple *DAM* genes have lost during evolution. Similar functional diversification of *SVP*-like genes during evolution was identified in kiwifruit, in which *AdSVP1* and *AdSVP3* complemented the early-flowering phenotype of the *svp-41* mutant but not *AdSVP2*, which is involved in delayed budbreak of lateral floral buds [87]. Moreover, whereas the central role of SVP TFs as a part of transcriptional complexes that regulate developmental processes seems conserved across taxa, DAM TFs might have evolved to regulate dormancy-specific functions in Rosaceae (as discussed in the previous section). This dormancy-related function is believed to be similar to the role that FLC plays in the control of flowering mediated by vernalization in Arabidopsis [88,89]. Therefore, it would not be surprising that the dormancy cycle is controlled by FLC-like TFs in trees. This seems not to be the case at least for the FLC-like TF studied in this work, as its expression pattern suggests a role related to ecodormancy and independent of cold requirement (Figure 2D, [5,46]). Moreover, during the evolution of the Rosaceae, *SVP*-like genes were expanded and several *FLC*-like genes were lost [83], favorizing the neofunctionalization of some *SVP*-like genes to take dormancy-related roles, and thus originating the MdDAM TFs. Indeed, in some non-Brassicaceae species, the FLC-like function is performed by distinct MADS TFs. For example, in wheat (*Triticum aestivum*), vernalization is regulated by *VERNALIZATION1* (*VRN1*), which encodes a MADS TF similar to the Arabidopsis APETALA1 TF [90].

After a process of neofunctionalization, a subsequent subfunctionalization event within the *DAM* genes could have provided more flexibility to the GRNs controlling the dormancy cycle in fruit trees. We believe that this could be due to the differential expression patterns shown by the *DAM* genes, as previously proposed by [15,91] (Figure 2), but also by the combination of DAM TFs in transcriptional complexes with distinct DNA-binding preferences, as shown by our seq-DAP-seq data (Figure 3 and Supplemental Figures S7 and S10). The oligomerization between MADS TFs is facilitated by their K domain (Keratin-like domain, a predicted coiled-coil) [33]. Interestingly, Rosaceae SVP-like TFs are highly variable in the K domain, which present a large number of potential positive selection sites [33,83]. Therefore, evolutionary forces might have contributed not only to originate a dormancy-specific group of TFs but also to combinatorial complexes that modulate the progression of dormancy in Rosaceae species.

### Conclusion

Trees have developed bud dormancy as a mechanism of adaptation against low temperatures of winter. Temperate fruit trees have been domesticated since thousands of years to adapt their dormancy and flowering patterns to local climatic conditions. However, the rapid increase of temperatures that is being observed in several world’s regions due to global warming is threatening food security by affecting the dormancy cycle progression and thus, flowering time and synchronization. Breeding for these traits requires a detailed knowledge on the genetic and molecular mechanisms underlying their biological regulation. Here, we presented a comprehensive genomic study focused on a group of MADS TFs that are known to regulate the dormancy cycle in Rosaceae fruit trees. The artificial downregulation of *MdDAM* and *MdSVP* genes in transgenic apple trees leads to non-dormant phenotypes that are able to flower regardless of the environmental conditions [4,23]. Thus, we believe that the new understanding that we produced on how these MADS TF are organized in transcriptional complexes and the identification of their genome-wide target genes is instrumental for generating novel varieties better adapted to future climatic conditions.

### Methods

#### Plant material

‘Royal Gala’ trees grafted on ‘M9’ rootstocks were distributed in three sampling blocks of four plants each in an orchard located at SudExpé experimental station in Marsillargues, France. Three dormant bud meristems were harvested per plant in eight timepoints from November 11^th^ 2016 to February 27^th^ 2017. Samples were immediately frozen in liquid nitrogen in the field and stored at −80 °C until use. Once a week after January 2017, shoots (axillary branches of around 50 cm) containing 10 buds each were harvested in the field and the dormant stage of the buds was evaluated under forcing conditions (16 h light/8 h dark and 22 °C) using the Tabuenca’s test [92]. The accumulation of chilling hours (number of hours with temperatures below 7.2 °C) was monitored by daily recording the air temperature in field trials (Supplemental Figure S6).

#### Gene expression studies in apple

Total RNA was isolated using the Spectrum^™^ Plant Total RNA kit (Sigma-Aldrich), and DNase-treated using the TURBO DNA-free Kit (Ambion). The SuperScript^™^ III First-Strand Synthesis System (Thermo Fisher Scientific) was used for cDNA synthesis according to manufacturer’s instructions. Real-time PCR was performed using the LightCycler 480 instrument (Roche), and relative expression was calculated using the 2(^-ΔΔ^Ct) method as described in [93,94]. The RT-qPCR primers are listed in Supplemental Table S1. *MdMDH* and *MdWD40* were used as reference genes [95].

#### *pENTR* vector construction

The complete coding sequences of *MdDAM1, MdDAM2, MdDAM4, MdDAMb, MdSVPa, MdSVPb, MdFLC, SVP* and *Venus* were amplified with high fidelity enzymes (Merck) and cloned into *pDONR201* and *pDONR207* vectors [96] using the Gateway^®^ BP Clonase^™^ II Enzyme Mix (Thermo Fisher Scientific). Truncated versions retaining the MADS (MADS-box), I (intervening), K (K-box) and a minimal C-terminal region were generated for all seven apple genes using gene-specific primers (Supplemental Table S1), and the obtained sequences were cloned into *pDONR201*. The CDS of *MdDAM1, MdDAM4, MdFLC* and *MdSVPa* without the stop codon were independently amplified and cloned into *pDONR207*. These constructs were then recombined into the *pBEACON-GR* vector kindly provided by Dr. Gloria Coruzzi (New York University, USA) using the Gateway^®^ LR Clonase^™^ II Enzyme Mix (Thermo Fisher Scientific). Genes fused to GR were amplified and cloned into the *pDONR207* vector. The vectors resulting from each cloning step were confirmed by sequencing.

#### Complementation assay in Arabidopsis

The CDS of *MdDAM1, MdDAM2, MdDAM4, MdDAMb, MdSVPa, MdSVPb, SVP* and *Venus* were independently introduced into a modified *pGreen0229* Gateway vector (*pYB187*, kindly provided by Dr. Youbong Hyun, MPIPZ, Cologne) [97] in which the p*35S* promoter was replaced by the Arabidopsis *SVP* promoter. To this end, overlapping primers were designed to amplify a region of 3 kb upstream of the TSS of *SVP* with complementary borders to the *pYB187* vector (Supplemental Table S1). In parallel, PCR reactions were performed to amplify the *pYB187* plasmids carrying the cloned CDS sequences. Overlapping fragments were assembled by Polymerase Incomplete Primer Extension [98]. All constructs were introduced into Agrobacterium strain GV3101 [99] and Arabidopsis *svp-41* mutant plants [45] were transformed using the floral dip method [100]. Several T1 independent lines were BASTA selected for each construct (Supplemental Figure S1), lines showing a Mendelian segregation (3:1) in the next generation were followed, and two to three homozygous single copy T3 lines from independent T1 lines were used in further studies. Seeds were stratified on soil for 7 days in the dark at 4 °C. Plants were grown under controlled environmental conditions at 22 °C in LDs (16 h light/8 h dark). Arabidopsis Columbia-0 (Col-0) was used as WT. Flowering time was scored by counting total leaf number (cauline and rosette leaves) of at least 10 plants per genotype. Number of days from germination to bolting (elongation of the first internode around 0.5 cm) and to the opening of the first flower were recorded. Total RNA was isolated from 7-day-old manually dissected leaves and shoot apices (containing a segment of the apical stem, SAM and young leaves) using the RNeasy mini kit (QIAGEN). DNase treatment, cDNA synthesis and RT-qPCR analysis were performed as previously described. *PP2A* was used as internal control.

#### Yeast two-hybrid

Full-length and truncated versions of the apple MADS genes were recombined into *pDEST22* (Activation Domain – AD) and *pDEST32* (DNA-Binding Domain – BD) vectors (Invitrogen). The protein–protein interactions to be tested were transformed into yeast PJ69-4A strain following the Frozen-EZ Yeast Transformation II^™^ (Zymo Research) protocol. Yeast selection was initially carried out in SD plates lacking Leu and Trp (–L–W). Three to five colonies were randomly selected, mixed and grown on SD plates lacking –L–W or Leu, Trp and His (–L–W–H) supplemented or not with different concentrations of 3-amino-1,2,4-triazole (3-AT). Yeast were grown at 30 °C for six days.

#### Tobacco co-immunoprecipitation

Full-length gene versions were fused to N-terminal 5xMyC or GFP by Gateway recombination into the pAM backbone [101]. A modified pEarleyGate301 [102] Gateway vector (kindly provided by Dr. Diarmuid O’Maoileidigh, MPIPZ, Cologne), in which the *att* borders, the *HA* and the *OCS* sites were replaced by the *Pro35Sx2::NLS-Venus* cassette, and *Pro35S::5xMyC-AtHB33* were used as controls. Binary vectors were transformed into Agrobacteria as previously described. Different protein combinations were transiently co-infiltrated in *N*. *benthamiana* leaves as described elsewhere [40]. For each combination, two tobacco plants were used, and three leaves per plant were infiltrated. Tobacco plants were incubated at room temperature for 3 days, and leaves were harvested and frozen in liquid nitrogen. Tobacco leaves were ground in liquid nitrogen and resuspended in extraction buffer (50 mM Tris–HCl pH 7.5, 150 mM NaCl, 10% glycerol, 2 mM EDTA pH 8 with HCl, 5 mM DTT, 0.2% Triton, plant protease inhibitor cocktail (Sigma)). 35 ug of total protein was mixed with 2X Laemmli Buffer to be used as input. Protein concentration was normalized to 0.7 mg of total protein, and the supernatant was incubated with the GFP-trap^®^_A kit (Chromotek) to immunoprecipitate GFP-fused proteins. Beads were washed three times, resuspended in 2X Laemmli Buffer and boiled at 96 °C for 10 min. Western blot analysis was performed to detect immunoprecipitated proteins using anti-GFP monoclonal antibody (ab290 from Abcam) and co-immunoprecipitated with anti-MyC monoclonal antibody (9E1 from Chromotek) as described [40]. Chemiluminescence detection of the proteins was done using the ChemiDoc MP Imager (Bio-Rad).

#### Confocal microscopy

To visualize Venus expression in Arabidopsis leaves (7-day-old) and shoot meristems (10-day-old), the method of [103] was used with the modifications proposed by [104]. Samples were imaged by confocal laser scanning microscopy (SP8; Leica) using settings optimized to visualize Venus fluorescence (laser wavelength, OPSL 514 nm; detection wavelength, 521 to 531 nm) and Renaissance 2200 (laser wavelength, Diode 405 nm; detection wavelength, 424 to 478 nm). To visualize GFP expression in tobacco leaves, the abaxial side of leaves were imaged after 3 days of incubation. The SP8 confocal microscope was set up to visualize GFP fluorescence (laser wavelength, OPSL 488 nm; detection wavelength 492 to 517 nm), and the bright field.

#### Protein production and purification for EMSA and seq-DAP-seq experiments

The full-length CDS sequences of *MdDAM1, MdDAM4, MdDAMb, MdFLC* and *MdSVPa* were amplified without the stop codon and the restriction sites of *EcoRI* and *SalI* were added to their 5’ and 3’, respectively (Supplemental Table S1). Each gene was amplified and cloned into the double *EcoRI-* and *SalI*-digested *XLp34* and *XLp39* vectors [43], fusing the genes to 3xFLAG and 5xMyC, respectively. In pairs, tagged proteins were simultaneously produced *in vitro* using the TNT^®^ SP6 High-Yield Wheat Germ Protein Expression System (Promega). Protein complexes were purified using anti-FLAG magnetic beads (Merck Millipore), and Western blots were performed as described above. For EMSA and seq-DAP-seq assays, protein complexes were co-produced *in vitro* and sequential pull-downs were carried out using anti-FLAG and anti-MyC (Thermo Scientific) beads [43]. EMSA was performed as described [105], and the DNA probe (10 nM) consisted of a 103 bp fragment of the Arabidopsis *SEP3* promoter containing two CArG boxes. The Arabidopsis AG–SEP3 protein complex was used as a positive control. Seq-DAP-seq was performed essentially as described [43]. DNA libraries were prepared without any PCR amplification step in order to maintain the natural DNA methylation state of the biological samples. Thirteen seq-DAP-seq libraries were generated using apple genomic DNA extracted from Gala endodormant buds, three replicates for MdDAM1–MdSVPa, MdDAM4–MdSVPa, MdFLC–MdSVPa and MdSVPa–MdSVPa, and the input DNA as a control. Libraries with different barcodes (NEBNext^®^ Multiplex Oligos for Illumina^®^) were pooled with equal molarity, and sequenced on Illumina HiSeq (Genewiz, https://www.genewiz.com/) with specification of paired-end sequencing of 150 bp. Each library obtained between 40 and 60 million reads (730 million of total reads).

#### Calli transformation and RNA-seq

The genes fused to GR previously cloned in *pENTR* vectors were recombined into the binary vector *pCamway35S* [106], which carries the constitutive promoter *CaMV35S* driving transgene expression. The expression vectors were used in the transformation of *A. tumefaciens* as described above. *In vitro* cuttings of apple cultivar Gala were subcultured, and apple leaf transformation was carried out to produce transformed calli as described elsewhere [78]. Positive transformed calli were selected in media supplemented with antibiotics and the observation of GFP fluorescence. Six months after leaf transformation, 30 transformed calli were obtained for each construct. Transformed calli were pretreated with 40 μM CHX (Sigma-Aldrich) for 30 min, rinsed with distilled water, and treated with 10 μM DEX (Sigma-Aldrich) or mock (ethanol). After 1 h, samples were rinsed with distilled water, incubated at room temperature for 7 h and then were frozen in liquid nitrogen and stored at −80 °C. RNA isolation was performed as previously described. Samples in triplicate were used to generate 24 Illumina Truseq Stranded mRNA libraries that were sequenced at the GeT Platform (https://get.genotoul.fr/en/) using an Illumina HiSeq 3000 with specification of paired-end sequencing of 150 bp. Each library obtained between 20 and 40 million reads (670 million of total reads).

#### Bioinformatic analyses

##### Read cleaning and mapping of seq-DAP-seq assays

Raw seq-DAP-seq reads were pre-processed by removing potential sequence adapters with Cutadapt [107] and trimming of low-quality bases (Q < 15) at the ends using Trimmomatic [108]. All pre-processed reads with final lengths smaller than 50 bases were discarded. Cleaned reads were mapped to the apple double-haploid genome version 1.1 [56] using BWA [109] with default settings. Raw BWA alignments were filtered by only keeping alignment pairs with mapping quality of at least 20 and by removing secondary alignments.

##### Peak calling and annotation

Raw seq-DAP-seq peaks were individually called together with a single control sample (input) using the program MACS2 [110] (parameters: -p 0.05 -g 550601767). Individual peak sets were called for all three replicates of the four different complexes. Reproducibility between the replicate peak sets was assessed using the irreproducible discovery rate (IDR) of 0.01 with the framework available from the BioConda channel of the conda package manager [111]. The final peak sets were generated by merging overlapping peaks that passed the IDR assessment. Peaks were annotated to apple genes (3 kb up- and 1 kb downstream) using the Bioconductor R package ChIPpeakAnno [112].

##### Peak position analysis

The preferred location of peaks relative to genes was analyzed using a previously described approach [113]. Briefly, for every peak annotated to a gene, the distance between the peak center to the TSS or TES of the respective gene was calculated depending on whether the peak was located upstream or downstream of it, respectively. For those peaks residing within the gene body, a normalized distance was calculated in relation to the TSS. The normalized distance was obtained using a gene-specific length factor that sets the gene length to 2,000 nucleotides. The observed peak location distribution was compared to that of 1,000 random peak sets consisting of the exact number of peaks with the same length as the observed peak set.

##### Identification of enriched motifs

*De novo* motif enrichment analysis was performed using MEME-chip [114]. For that, the central 100 nucleotides of the peaks were used for motif discovery (parameters: meme-chip-meme-minw 8 -meme-maxw 15 - meme-nmotifs 15 -meme-mod anr -db JASPAR2018_CORE_plants_non-redundant_pfms_meme.txt -centrimo -local -centrimo-ethresh 1), provided with binding sites of plant-specific transcription factors obtained from the JASPAR webpage (http://jaspar.genereg.net). The occurrences of predicted motifs were extracted from the corresponding FIMO output produced by the MEME-ChIP suite. Complementally, a custom C++ code was written to manually search the peaks for the presence of a previously described degenerated CArG-box motif, MYHWAWWWRGWWW [88,115].

##### RNA-seq analyses

The RNA-seq reads of the GR DEX-inducible assays were pre-processed as described above (see “Read cleaning and mapping of seq-DAP-seq assays”). Cleaned reads were mapped to the apple genome using TopHat2 [116] (parameters: -i 50 -I 10000 -N 5 –read-edit-dist 5 –read-gap-length 2). The apple genome annotation was used for intron hints. The number of reads mapped to each gene was determined using the HTSeq count program [117]. Differential gene expression analyses were carried out using the DESeq2 R package [118]. The publicly available transcriptome samples PRJNA374502 (dataset A [4]) and PRJDB6779 (dataset B [5]) were downloaded from the NCBI sequence read archive and quantified using the Salmon program [119]. Differential gene expression analyses were performed as previously described.

##### Gene clustering

FPKM gene expression values for dataset A [4] were obtained using the fpkm function of DESeq2 [118]. The expression values were transformed into the z-scale and clustered using the scale and hclust functions of R [120], respectively. The dendrograms from the heatmaps generated from the z-transformed expression data were visually inspected and a cutoff value was arbitrarily chosen in order to obtain the gene clusters. The expression data for each cluster was summarized by the eigen gene, which was obtained using the moduleEigengenes function of the R package WGCNA [121].

##### GO enrichment tests

A BLAST [122] similarity search (parameters: -F F -e 1e-10 -b 1 -v 1) was performed using the apple proteins as a query against the Arabidopsis TAIR10 proteome (www.arabidopsis.org). The apple proteins/genes inherited the GO term annotation of their best Arabidopsis counterpart.

The ‘plant GO slim’ subset was obtained from the GO website (http://current.geneontology.org/ontology/subsets/goslim_plant.obo). All enrichment tests using the ‘plant GO slim’ annotations were performed in MATLAB. By using the geneont class from the MATLAB Bioinformatics toolbox, genes were annotated with a particular GO slim term if one of its originally blast-based GO terms corresponds to or is a descendant of that GO slim category. The GO term enrichment in a gene set was assessed using the hypergeometric test. Raw P-values were adjusted for multiple testing using the Benjamin Hochberg method [123]. For the enrichment analysis of the seq-DAP-seq targets of the four MADS complexes, the background consisted of all genes in the apple genome with at least one GO slim annotation. The test sets encompassed the target genes of each MADS complex having the specific GO slim terms to be tested. The results were displayed as heatmaps with a grayscale reflecting the P-value. For the enrichment analysis using expression data from the calli assay, the background was determined as the set of genes tested by DESeq2 (in a MADS TF-dependent manner) and assigned to at least one GO slim term. The test sets consisting of GO-annotated DEGs were then separated into up- or downregulated gene sets, and both sets were individually tested for GO slim term enrichment. The results were plotted as a heatmap in which an orange or blue gradient was assigned in the event that the upregulated or the downregulated set, respectively, was enriched (P ≤ 0.05). In case both sets were enriched, the set with the best P-value was kept.

The GO term enrichment analysis for the gene clusters obtained from dataset A [4] was performed using the Cytoscape plugin BiNGO [124]. For these tests, the background consisted of all GO annotated apple genes. The test sets consisted of the genes within a particular cluster having the GO term being tested.

## Statistical analysis

Statistical analysis using Kruskal-Wallis one-way ANOVA, ANOVA and t tests were conducted using Prism 5.0a (GraphPad). Fisher and hypergeometric tests were conducted using R [120].

## Declarations

### Ethics approval and consent to participate

Not applicable.

### Consent for publication

Not applicable.

### Availability of data and materials

The datasets generated and/or analysed during the current study are available in the NCBI SRA repository, under the accession code PRJNA698061.

### Authors’ contributions

V.S.F, E.S, X.L, J.E, I.F and V.H performed the experiments, V.S.F and E.S analyzed the data, V.S.F, L.F.R, C.Z, G.C, E.C and F.A conceived and designed the experiments, V.S.F, E.S and F.A wrote the manuscript. All the authors approved the final version.

## Acknowledgements

We thank Dr. Gloria Coruzzi (New York University, New York, New York, USA) and Dr. Julie Leclercq (AGAP, University of Montpellier, Montpellier, France) for generously providing materials. We also thank to Dr. Virginia Fernandez and Dr. Eva Madrid Herrero (MPIPZ, Cologne, Germany) for helping with the experimental setup, and François Parcy for hosting V.S.F in his laboratory for a short-term stay.

## Legend to supplemental information

**Supplemental Figure S1. Flowering-time distributions shown as total leaf number of T1 populations transformed with the different apple *DAM*-like, *SVP*-like or control genes.** X-axis shows rosette numbers of leaves of individual plants, and y-axis shows the numbers of individual plants. The red square represents the median for each gene population.

**Supplemental Figure S2. Expression pattern of *pSVP::Venus* lines.** Confocal analysis of expanded leaves (7-day-old plants) or dissected apices (10-day-old plants) of *pSVP::Venus* lines.

**Supplemental Figure S3. Protein–protein interactions among apple DAM-, SVP- and FLC-like proteins. A)** Yeast two-hybrid assay using full-length apple proteins. MdDAM1, MdDAM2 and MdDAM4 showed strong autoactivation even in the drop-out media supplemented with 5 mM of 3AT. To circumvent this issue, the C-terminal region of all proteins was removed, as it is known that this region is responsible for the generation of autoactivation in MADS-domain proteins. **B)** Yeast two-hybrid assays using truncated protein versions. Interactions were tested in both directions, unless a positive interaction was identified before. In this case, the other direction was not tested (blank spaces with nt – not tested). Proteins that did not produce autoactivation were not evaluated in the drop-out media supplemented with 3AT. The negative controls are representative pictures of several independent controls that were used in this assay.

**Supplemental Figure S4. Subcellular localization of apple DAM-, SVP- and FLC-like proteins.** MdDAM1 **(A)**, MdDAM2 **(B)**, MdDAM4 **(C)**, MdDAMb **(D)**, MdSVPa **(E)**, MdSVPb **(F)** and MdFLC **(G)** were translationally fused to GFP and *Nicotiana benthamiana* leaves were agroinfiltrated and observed in a confocal microscope after 3 days of incubation. **H)** VENUS fused to a nuclear localization signal (NLS). Scale bar 100 μm.

**Supplemental Figure S5. Flowering time under LD of Arabidopsis F1 plants in comparison to WT and homozygous lines. A)** Total leaf number (including both cauline and rosette leaves) was scored prior to flowering. **B)** Number of days from germination to bolting (elongation of the first internode around 0.5 cm). **C)** Number of days from germination to the opening of the first flower. The box extends from the 25th to 75th percentiles, the line in the middle is plotted at the median, and the whiskers are drawn down to the 10th and up to the 90th percentile. The outliers below and above the whiskers are drawn as individual points. One-way ANOVA followed by Tukey’s test was used for the statistical tests. Letters shared in common between the genotypes indicate no significant differences (for P ≤ 0.05).

**Supplemental Figure S6. Chilling hours accumulated during the dormancy cycle.** One chilling hour was considered as one hour with temperatures below 7.2°C. Red arrows represent sampling points for RT-qPCR studies.

**Supplemental Figure S7. Complementary information about the seq-DAP-seq assays. A)** *In vitro* co-immunoprecipitation of proteins. MdDAM1, MdDAM4, MdFLC and MdSVPa were translationally fused to FLAG, whereas MdDAM1, MdDAM4, MdDAMb, MdFLC and MdSVPa were translationally fused to MyC. Protein fusions were synthesized *in vitro* and tested in pairs. The input was composed of total proteins recovered before immunoprecipitation. FLAG-fused proteins were immunoprecipitated using anti-FLAG beads and immunoblotted using anti-FLAG or anti-MyC antibody. **B)** Logos of the enriched CArG-box motifs identified with the degenerated CArG-box strategy (see Methods). **C)** GO term enrichment analysis of target genes containing a CArG-box for each apple transcriptional complex. For data visualization, the best P-value was transformed using log. Note that all P-values higher than 0.05 were replaced by zero (white boxes). **D)** DNA-binding profiles of the four complexes and the control (input) to the locus region of *MdDAM1, MdDAMb* and *MdFLC*. Horizontal bars below the plots represent the position of the peak regions. The color code between the plots and the bars is preserved. The Integrated Genome Browser (IGB) was used for visualization.

**Supplemental Figure S8. Complementary information about the GR DEX-inducible assays. A)** Dendrogram illustrating the distribution of the RNA-seq replicates in the GR DEX-inducible assay performed in transgenic apple calli. **B)** Venn diagram illustrating common DEGs between the four TFs using the GR DEX-inducible system. **C)** GO term enrichment analysis of genes identified in transgenic apple calli after 8 hours of DEX induction. Enrichment tests were performed separately for up- and downregulated gene sets and only the best P-value (smallest) was kept. For data visualization, the best P-value was transformed using -log or log when it belonged to the up (orange gradient) or downregulated (blue gradient) gene set, respectively. Note that all P-values higher than 0.05 were replaced by zero (white boxes).

**Supplemental Figure S9. Expression analysis of *MdDAM1, MdDAM4, MdFLC* and *MdSVPa* in two public RNA-seq during apple dormancy. A)** Time-course expression analysis of apple buds harvested from field-grown ‘Golden delicious’ trees according to Moser et al. [4]. **B)** Expression analysis of apple bud samples harvested from ‘Fuji’ trees in the field and exposed to controlled chilling conditions according to Takeuchi et al. [5]. **C)** Venn diagram illustrating the overlap between dormancy-related DEGs. Only genes present in both datasets were considered in this analysis. The obtained P-value (hypergeometric test) for the intersection between datasets was equal to 0.

**Supplemental Figure S10. MADS transcriptional complexes regulate several dormancy-related genes. A)** DNA-binding profiles of four MADS transcriptional complexes and the control (input) to the locus region of several dormancy-related genes. **B)** DNA-binding profiles of four MADS transcriptional complexes and the control (input) to the locus region of *MdSOC1a*. Time-course expression analysis of apple buds harvested from field-grown ‘Golden delicious’ trees according to Moser et al. [4]. The Integrated Genome Browser (IGB) was used for visualization. The arrows represent induction or repression according to the expression data obtained in the calli assay for genes belonging to the high-confidence list of target genes (Supplemental Data S4).

**Supplemental Table S1.** List of primers employed in this work.

**Supplemental Data S1.** List of seq-DAP-seq peaks for the four MADS transcriptional complexes.

**Supplemental Data S2.** List of seq-DAP-seq peaks containing at least one CArG-box (see Methods) for the four MADS transcriptional complexes.

**Supplemental Data S3.** RNA-seq result of the GR DEX-inducible assay in apple calli for four MADS TFs.

**Supplemental Data S4.** List of high-confidence targets of four MADS transcriptional complexes.

**Supplemental Data S5.** Heatmap summarizing the gene expression during dormancy of the high-confidence targets of MdDAM1–MdSVPa, MdDAM4–MdSVPa and MdFLC–MdSVPa complexes.

**Supplemental Data S6.** Heatmap summarizing the gene expression during dormancy of the high-confidence targets of MdDAM1–MdSVPa complex.

**Supplemental Data S7.** Heatmap summarizing the gene expression during dormancy of the high-confidence targets of MdDAM4–MdSVPa complex.

**Supplemental Data S8.** Heatmap summarizing the gene expression during dormancy of the high-confidence targets of MdFLC–MdSVPa complex.

**Supplemental Data S9.** Enrichment analysis on GO terms for biological processes over the clusters formed during dormancy for the target genes of MdDAM1–MdSVPa, MdDAM4–MdSVPa, and MdFLC–MdSVPa complexes.

**Supplemental Data S10.** List of genes differentially regulated during dormancy is dataset A [4] and dataset B [5].

**Supplemental Data S11.** List containing 231 dormancy-related DEGs that are high-confidence targets of MdDAM1–MdSVPa, MdDAM4–MdSVPa, and/or MdFLC–MdSVPa complexes.

